# An extended admixture pulse model reveals the limitations to Human-Neandertal introgression dating

**DOI:** 10.1101/2021.04.04.438357

**Authors:** Leonardo N. M. Iasi, Harald Ringbauer, Benjamin M. Peter

## Abstract

Neandertal DNA makes up 2-3% of the genomes of all non-African individuals. The patterns of Neandertal ancestry in modern humans have been used to estimate that this is the result of gene flow that occurred during the expansion of modern humans into Eurasia, but the precise dates of this event remain largely unknown. Here, we introduce an extended admixture pulse model that allows joint estimation of the timing and duration of gene flow. This model leads to simple expressions for both the admixture segment distribution and the decay curve of ancestry linkage disequilibrium, and we show that these two statistics are closely related. In simulations, we find that estimates of the mean time of admixture are largely robust to details in gene flow models, but that the duration of the gene flow can only be recovered if gene flow is very recent and the exact recombination map is known. These results imply that gene flow from Neandertals into modern humans could have happened over hundreds of generations. Ancient genomes from the time around the admixture event are thus likely required to resolve the question when, where, and for how long humans and Neandertals interacted.

## Introduction

The sequencing of Neandertal (Green et al., 2010; Prüfer et al., 2013, 2017; Mafessoni et al., 2020) and Denisovan genomes (Reich et al., 2010; Meyer et al., 2012) revealed that modern humans outside of Africa interacted, and received genes from these archaic hominins (Vernot and Akey, 2014; Fu et al., 2014; Sankararaman et al., 2014; Fu et al., 2015; Malaspinas et al., 2016; Sankararaman et al., 2016; Vernot et al., 2016). There are two major lines of evidence: First, Neandertals are genome-wide more similar to non-Africans than to Africans (Green et al., 2010). This shift can be explained by 2-4% of admixture from Neandertals into non-Africans (Green et al., 2010; Prüfer et al., 2013). Similarly, East Asians, Southeast Asians and Papuans are more similar to Denisovans than other human groups, which is likely because of gene flow from Denisovans (Meyer et al., 2012).

As a second line of evidence, all non-Africans carry genomic segments that are very similar to the sequenced archaic genomes. As these putative *admixture segments* are up to several hundred kilobases (kb) long, it is unlikely that they were inherited from a common ancestor that predates the split of modern and archaic humans (Sankararaman et al., 2014; Vernot and Akey, 2014). Rather, they entered the modern human populations through later gene flow (Sankararaman et al., 2012, 2014; Vernot and Akey, 2014; Sankararaman et al., 2016; Vernot et al., 2016).

However, substantial uncertainty remains about when, where, and over which period of time this gene flow happened. A better understanding of the location and timing of the gene flow would allow us to place constraints on the timing of movements of early humans, and the population genetic consequences of their interactions.

Archaeological evidence puts some temporal boundaries on the times when Neandertals and modern humans might have interacted. The earliest currently known modern human remains outside of Africa are dated to around 188 thousand years ago (kya) (Hershkovitz et al., 2018; Stringer and Galway-Witham, 2018) and the latest Neandertals are suggested to have lived between 37 kya and 39 kya old (Higham et al., 2014; Zilhão et al., 2017). Thus the time window where Neandertals and modern humans might have been in the same area stretches over more than 140,000 years. However, there is less direct evidence of modern humans and Neandertals in the same geographical location at the same time. In Europe, for example, Neandertals and modern humans likely overlapped only for less than 10,000 years (Bard et al., 2020).

### Genetic Dating of Gene Flow

A common approach to learn about admixture dates from genetic data uses a *recombination clock* model: Conceptually, admixture segments are the result of the introduced chromosomes being broken down by recombination. The first generation offspring of an archaic and a modern human parent will have one whole chromosome each of either ancestry. Thus, the genomic markers in these individuals are in full ancestry linkage disequilibrium (ALD); all archaic variants are present on one DNA molecule, and all modern human variants on the other one.

If this individual has offspring in a largely modern human population, in each generation meiotic recombination will reshuffle the chromosomes, progressively breaking down the ancestral chromosome down into shorter segments of archaic ancestry (Falush et al., 2003; Gravel, 2012; Liang and Nielsen, 2014), and ALD similarly decreases with each generation after gene flow (Chakraborty and Weiss, 1988; Stephens et al., 1994; Wall, 2000).

This inverse relationship between admixture time and either segment length or ALD is commonly used to infer the timing of gene flow (Pool and Nielsen, 2009; Moorjani et al., 2011; Pugach et al., 2011; Gravel, 2012; Sankararaman et al., 2012; Loh et al., 2013; Hellenthal et al., 2014; Liang and Nielsen, 2014; Sankararaman et al., 2016; Pugach et al., 2018; Jacobs et al., 2019). Most commonly, it is assumed that gene flow occurs over a very short duration, referred to as an *admixture pulse*, which is typically modelled as a single generation of gene flow (e.g Moorjani et al., 2011). This model has the advantage that both the length distribution of admixture segments and the decay of ALD with distance will follow an exponential distribution, whose parameter is directly informative about the time of gene flow (Pool and Nielsen, 2009; Gravel, 2012; Liang and Nielsen, 2014).

In segment-based approaches, dating starts by identifying all admixture segments, which can be done using a variety of methods (Seguin-Orlando et al., 2014; Sankararaman et al., 2016; Vernot et al., 2016; Racimo et al., 2017; Skov et al., 2018). The length distribution of inferred segments is then used as a summary for dating when gene flow happened.

Alternatively, ALD-based methods use linkage disequilibrium (LD) patterns, without explicitly inferring the segments (Chimusa et al., 2018) (Figure 1 B). Instead, admixture dates are estimated by fitting a decay curve of pairwise LD as a function of genetic distance, implicitly summing over all compatible segment lengths (Moorjani et al., 2011; Loh et al., 2013).

**Figure 1:**
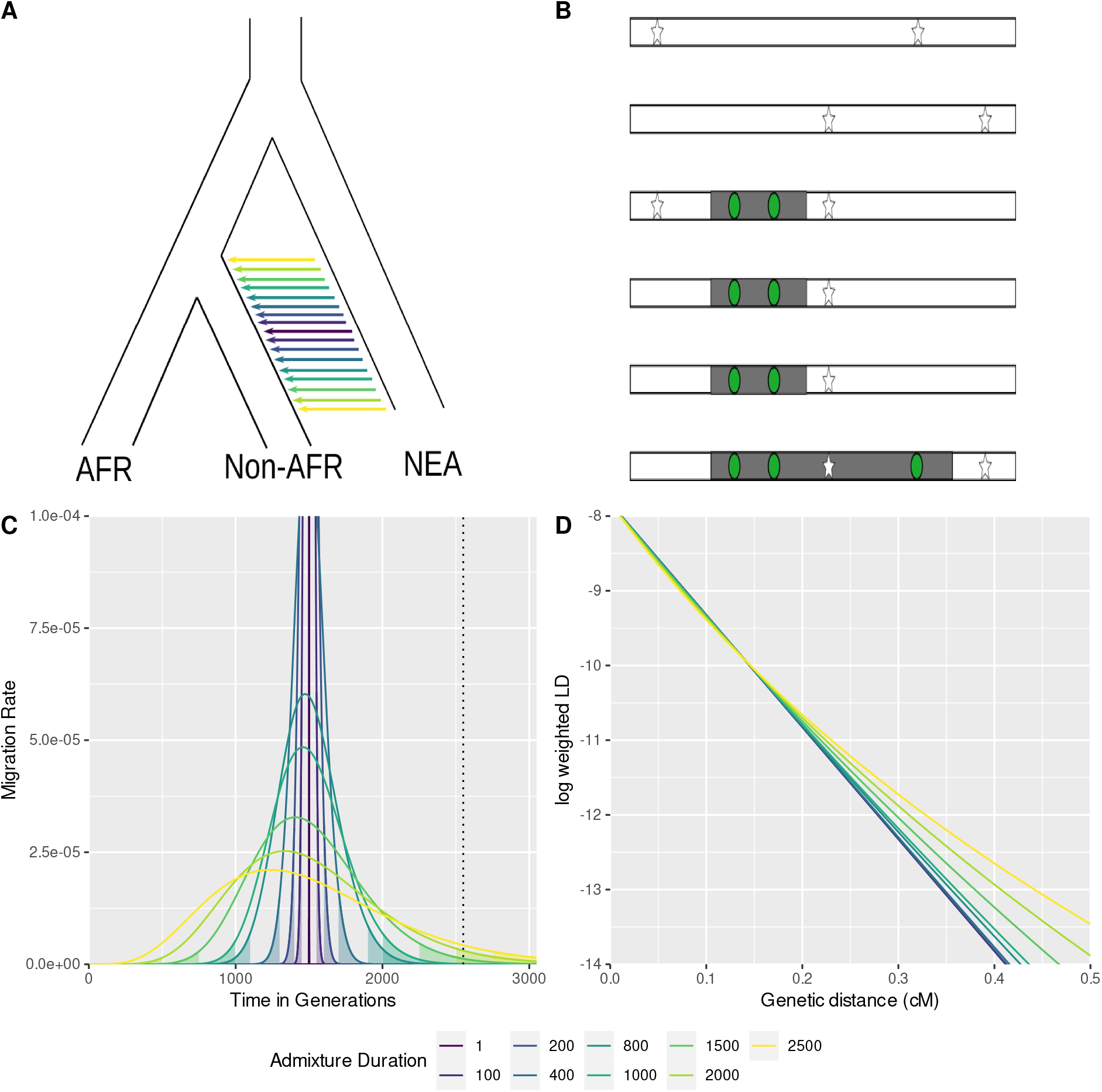
A) Neandertal introgression into non-Africans with a multitude of potential admixture durations. B) The time and duration of admixture results in different length distributions of introgressed chromosomal segments (grey) containing Neandertal variants (green circles) in high LD to each other compared to the background (human variants white stars). The ALD approach estimates linkage between the introgressed variants (green circles), whereas the haplotype approach tries to estimate the segment directly (grey area). C) Migration rate per generation modeled using the extended pulse model for different admixture durations (colored lines). The filled area under the curve indicates the boundaries of the discrete realization of the duration of gene flow *t_d_*. The dotted line indicates the oldest possible time of gene flow (as defined in the simulations). D) The expected LD decay under the extended pulse model.

### Neandertal Gene Flow Estimates

Using this approach, Sankararaman et al. (2012) dated the Neandertal-human admixture pulse to between 37–86 kya. Later, Moorjani et al. (2016) refined this date to 41 – 54 kya (*C.I*._95%_) using an updated method, a different marker ascertainment scheme and a refined genetic map for European populations. A date of 50 – 60 kya was obtained from the analysis of the genome of *Ust’-Ishim*, a 45,000-year-old modern human from western Siberia. The inferred Neandertal segments in *Ust’-Ishim* are substantially longer than those in present-day humans, which makes their detection easier, and adds further evidence that gene flow between Neandertals and modern humans has happened relatively recently before *Ust’-Ishim* lived (Fu et al., 2014). In addition, we have direct evidence of gene flow from early modern humans from Oase (Fu et al., 2015) and Bacho Kiro (Hajdinjak et al., 2021), dated to 40ky and 45ky, respectively. In genomes from both sites, segments of recent Neandertal ancestry less than ten generations before hint at admixture histories with late gene flow in Europe.

### Limitations of the Pulse Model

The admixture pulse model assumes that gene flow occurs over a short time period; however it is currently unclear how long a time could still be consistent with the data. This makes admixture time estimates hard to interpret, as more complex admixture scenarios might be masked, and so gene flow could have happened tens of thousands of years before or after the estimated admixture time.

That admixture histories are often complicated has been shown in the context of Denisovan introgression into modern humans, where at least two distinct admixture events into East Asians and Papuans were proposed (Browning et al., 2018; Jacobs et al., 2019; Choin et al., 2021). While the length distributions of admixture segments are similar between populations, there are differences in the genomic distribution of admixture segments, and their similarities to the sequenced high-coverage Denisovan (Browning et al., 2018; Massilani et al., 2020). In contrast, all Neandertal admixture segments are most similar to the Vindija Neandertal (Prüfer et al., 2017), but Neandertal ancestry is slightly higher in East Asians than Western Eurasians (Meyer et al., 2012; Wall et al., 2013; Kim and Lohmueller, 2015; Vernot and Akey, 2015; Villanea and Schraiber, 2019).

One way to refine admixture time estimates is to include two or more distinct admixture pulses. The distribution of admixture segment lengths will then be a mixture of the segments introduced from each event. This is especially useful if the events are very distinct in time, e.g. if one event is only a few generations back, and the other pulse occurred hundreds of generations ago (Fu et al., 2014, 2015). In this case, the admixture segments will be either very long if they are recent, or much shorter if they are older.

Zhou et al. (2017a) extended this model to continuous mixtures, using a polynomial function as a mixture density. However, they found that even for relatively short admixture events, the large number of parameters led to an underestimate of admixture duration (Zhou et al., 2017b).

### Extended Pulse Model

One drawback of these approaches is that they introduce a large number of parameters. Even a discrete mixture of two pulses requires at least three parameters (two pulse times and the relative magnitude of the two events) (Pickrell et al., 2014), and the more complex models require regularization schemes for fitting (Zhou et al., 2017b; Ralph and Coop, 2013).

Here, we propose an *extended admixture pulse* model (Figure 1 A) to estimate the duration of an admixture event. It only adds one additional parameter, reflecting the duration of gene flow, while retaining much of the mathematical simplicity present in the simple pulse model. The extended pulse model assumes that the migration rate over time is Gamma distributed, so that the length distribution of admixture segments has a closed form (Figure 1 C & D) with two parameters, the mean admixture time and duration.

Conceptually, identifying an extended pulse requires us to establish that the length distribution of admixture segments deviates from an exponential distribution. However, other sources of bias, such as the demography of the admixed population, the accuracy of the recombination map or details in the inference method parameters may also introduce similar biases. Thus, we have to carefully evaluate other potential sources of bias on whether they might lead to confounding signals. (Sankararaman et al., 2012; Fu et al., 2014; Moorjani et al., 2016).

Here, we first define the extended admixture pulse model and derive the resulting segment length and ALD distributions, and introduce inference schemes for either data. We then evaluate under which scenarios these two models can be distinguished. We show that power to distinguish these scenario is higher for more recent events and longer pulses, but that accurate inference requires high-quality data. Based on these results, we use data from European genomes (The 1000 Genomes Project Consortium, 2015) and find that for the case of Neandertal admixture, a simple pulses cannot be distinguished from continuous admixture over an extended period of time, and the data are consistent with a multitude of durations, up to several tens of thousands of years.

## New Approaches

In this section, we present the mathematical description of the admixture models we use in this paper, and introduce inference algorithms for estimating the admixture time and duration from both segment data and ALD.

### Admixture Models and Inference

We think of admixture as a series of “foreign” chromosomes introduced in a population (for a mechanistic model, see e.g. Pool and Nielsen (2009). Throughout, we assume that alleles evolve neutrally, and that recombination is independent of local ancestry. The simple pulse model assumes that all admixture happens in the same generation, (*i.e*. all chromosomes are introduced to the population at the same time). To extend this model, we allow chromosomes to enter at potentially many different time points, such that the migration rate at time *t* in the past is given by the function *m*(*t*) (Pool and Nielsen, 2009; Ni et al., 2016). For simplicity, we assume that the total amount of introgressed material 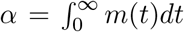 is small, so that segments do not interact, but we will discuss violations of this assumptions later. For archaic introgression, *α* ≈ 0.03, so this assumption is justified.

Over time, recombination splits up the introgressed genome into smaller pieces, while by the neutrality assumption the expected amount of total ancestry remains approximately the same. Thus, if we measure the size of chromosomes in recombination units, a chromosome of size *G* introduced at time *t* gives rise to an expected number of *tG* segments.

### Admixture Segment Lengths

We enumerate the admixture segments in a sample *i* = 1… *K*. We denote the length of the *i*-th segment as *L_i_* (measured in Morgan) and the time in the past when segment *i* entered the population as *T_i_* (measured in generations). We assume that the *L_i_* and *T_i_* are both realizations from more general distributions *L* and *T* that reflect the overall segment length and segment age distributions, respectively.

To relate *m*(*t*) to *T*, we need to take into account that older fragments had more time to split up (see e.g. Pool and Nielsen, 2009). Hence

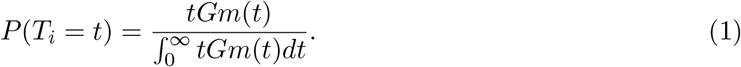

The denominator of the right-hand side term in equation 1 is the expected number of admixture segments, 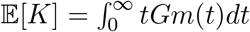.

Given *T_i_*, the segment length *L_i_* is exponentially distributed with rate parameter *t*:

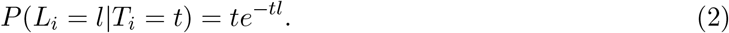

Integrating over *T* yields the unconditional distribution of admixture segment lengths:

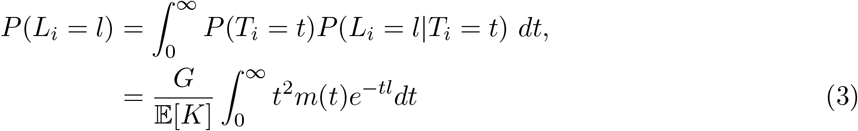

and we can think of *L* as an exponential mixture distribution with mixture density proportional to *tm*(*t*) (Ralph and Coop, 2013; Ni et al., 2016; Zhou et al., 2017a).

#### Ancestry Linkage Disequilibrium

Alternatively, the impact of gene flow is often characterized using ALD, particularly when accurate identification of archaic segments is difficult. We follow Loh et al. (2013) and note that the ALD from gene flow in a single event at time *t* generations in the past is

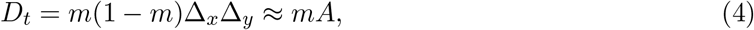

where *m* is the fraction of immigrants and Δ*_x_*, Δ*_y_* are the differences in allele frequencies between markers in the admixing populations. We assume that terms of the order of *m*^2^ can be ignored and that migration is low enough that changes in the allele frequencies in the admixing populations can also be neglected (i.e. *A* = Δ*_x_*Δ*_y_* remains a constant).

At a later generation *t*, the expected LD between two markers a distance *l* apart is

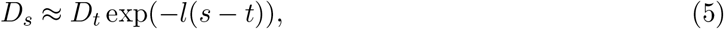

due to the decay of LD (e.g. Sankararaman et al., 2012). If the migration rate *m_f_* is a function of time, we can add up the LD introduced at each time *s* in the past and approximate *D* as

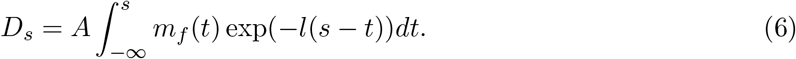

As we show in the Appendix (Formal Motivation for ALD), equation 6 satisfies the differential equation

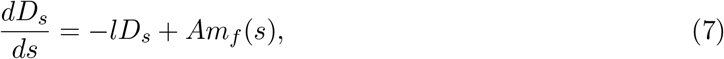

where the −*lD_s_*-term models the exponential decay of LD due to recombination, and the *Am_f_*(*s*)-term reflects the increase of LD due to admixture (Eq. 4).

To connect this equation more directly to the backward-in-time formulation used in the derivation of the admixture segment distribution, we set *s* = 0 and invert the flow of time, such that *m*(*t*) = *m_f_* (−*t*). We obtain

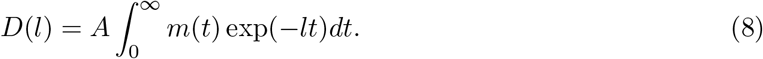

Thus, *D* can be interpreted as the tail function of an exponential mixture with mixture density *m*. Alternatively, the integral in equation 8 is also the (scaled) moment-generating function of *m* with argument −*l*.

The distribution of admixture segment lengths (equation 3) and the ALD function (equation 8) are closely related – in the Appendix (Connection Between Admixture Segment Length Distribution and ALD Function) we show that

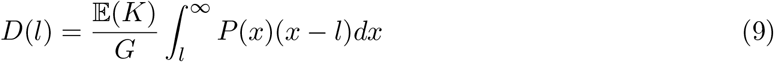

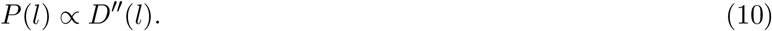

It follows that both functions uniquely determine each other. Consequently they contain identical information to estimate admixture dates.

Both for the segment and ALD models we use simplifying assumptions that ignore the effects of genetic drift, the recombination between introgressed segments and the replacement of older introgressed material. In the Appendix, we discuss these approximations and show that particularly the replacement of admixed material can be accommodated by replacing *m* with

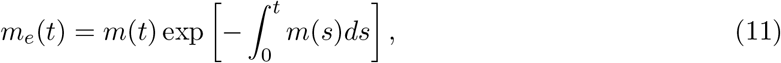

which can be interpreted as the probability of the event that migration happened at time *t*, and no more migration happened later on.

#### The Simple Pulse Model

Under the simple pulse model, all fragments enter the population at the same time *t_m_*, and *T* is a constant distribution. We can formalize this model by using a Dirac delta function which integrates to one if the integration interval includes *t_m_* and zero otherwise:

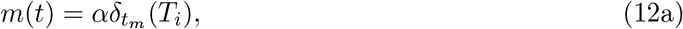

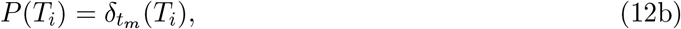

We obtain the exponential distribution of admixture fragments under this model (e.g. Moorjani et al., 2011):

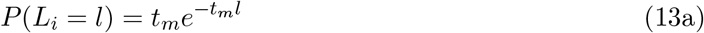

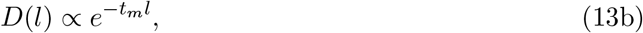

where here and in the remainder of this section we omit the constant term from *D*, which is not relevant for fitting the LD decay. The expected segment length under a simple pulse model is given by

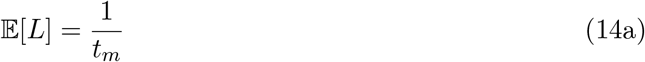

and the variance by

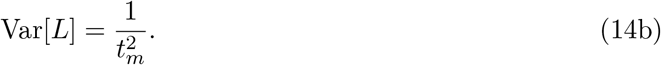

#### The Extended Pulse Model

For the new extended pulse model, we assume that the migration rate *m*(*t*) follows a rescaled Gamma distribution so that the total contribution of migrant alleles is *α*. It is convenient to parameterize the migration rate as 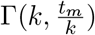. for *t* ≥ 0 and *k* ≥ 1.

Using this parameterization, the denominator of equation 1 is *t_m_αG* and

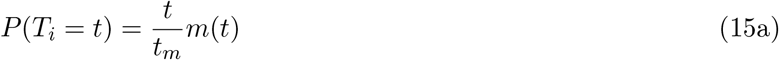

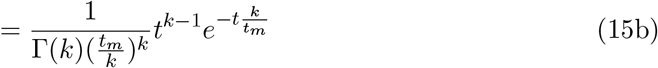

for *t* ≥ 0 and *k* ≥ 2, which is is the density of a 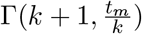-distribution with moments

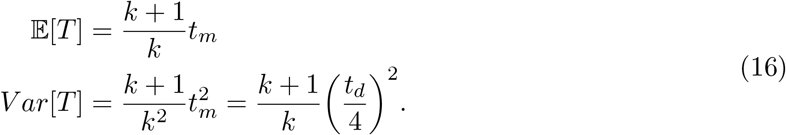

Here, we define the admixture duration 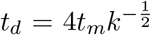, as a convenient measure for the duration of gene flow. If *k* is low, then *t_d_* will be large and gene flow extends over many generations. In contrast, if *k* is large, then *t_d_* ≈ 0 and we recover the simple pulse model (Figure 1 C & D).

The distribution of segment length is calculated by plugging equation 15b into equation 3 and integrating:

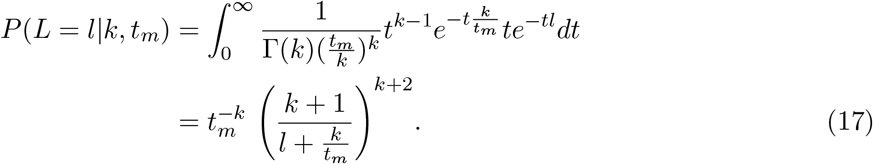

The distribution in equation 17 is known as a *Lomax* or *Pareto-II* distribution, which is a heavier-tailed relative of the Exponential distribution. Under the extended pulse model, the expected segment length will be the same as under the simple pulse model (Eq. 14a):

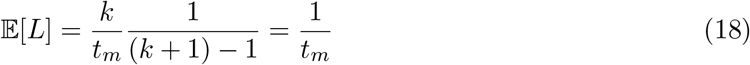

but the variance is larger:

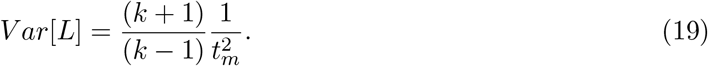

We obtain the ALD-function from equation using the moment-generating function of *m*(*t*):

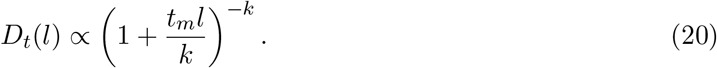

#### The Constant Migration Model

The simple pulse model can be thought of as the extreme case of the extended pulse model when *k* → ∞, i.e. the pulse gets infinitely short. In the other extreme the extended pulse model approaches a model of constant migration. In this case, the last migration event at a particular location is exponentially distributed with rate *m* (Eq. 11), which is a model considered by Pool and Nielsen (2009). Setting 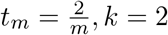, we obtain

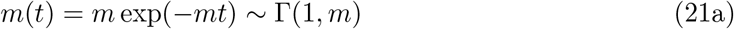

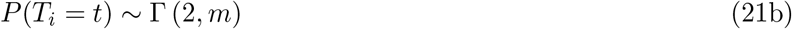

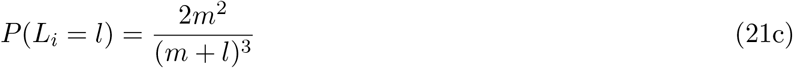

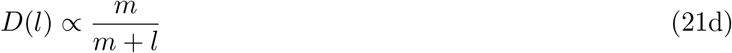

Equation 21c differs slightly from equation 6 in Pool and Nielsen (2009) because we approximate the expected number of segments with *n* = *Kt*, versus theirs *n* = 1 + *Kt* (however they converge to each other for large *Kt*).

#### Estimation of Admixture Times

For inference, either the admixture segment lengths or ALD can be used. Assuming the admixture segment lengths are known, equation 17 is the likelihood function and can be used for inference. For inference using ALD, we follow Moorjani et al. (2011) and use the decay of ALD with genetic distance as a statistic. Following Moorjani et al. (2016), we add an intercept *A* and a constant *c* modelling background LD:

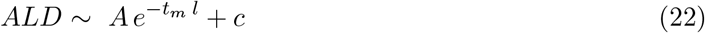

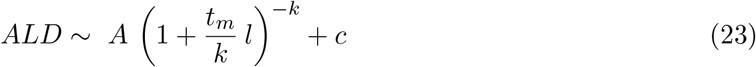

## Results

Here, we investigate under which scenarios we can distinguish the simple and extended pulse models, and when we can infer parameters under either model. We start with an idealized scenario of simulations under the model, and then continue with more realistic coalescent simulations using `msprime` (Kelleher et al., 2016).

### Power Analysis under the Model

In the easiest case, we assume that segments are known and we simulate directly under the model (Eq. 17) and evaluate under which conditions we can tell the two models apart using likelihood-ratio tests on the simulated segments. For this purpose, we compare two scenarios, one where gene flow happened 1,500 generations ago, which reflects Neandertal gene flow inferred from present-day individuals. In the second scenario, which reflects inference from ancient modern human data, the samples are taken 50 generations after gene flow ended. We vary pulse durations from one to 2,500 generations, and sample between 100 and 100,000 unique segments. As the simple pulse model is an edge case of the extended pulse model with *k* → ∞, standard likelihood theory does not apply, and we use empirical significance cutoffs (Kozubowski et al., 2008).

The resulting log-likelihood ratios are given in Figure 2. In general, we find that power to distinguish the model increases with pulse duration and the amount of data, and that it is easier to distinguish the models when gene flow had been more recent. For example, with 10,000 unique segments we need an event lasting around 1,000 generations before we are able to confidently distinguish an extended from a simple pulse (Figure 2) using present-day data. In contrast, by sampling closer to the admixture event we are able to distinguish an extended pulse already with a duration of 40-60 generations.

**Figure 2:**
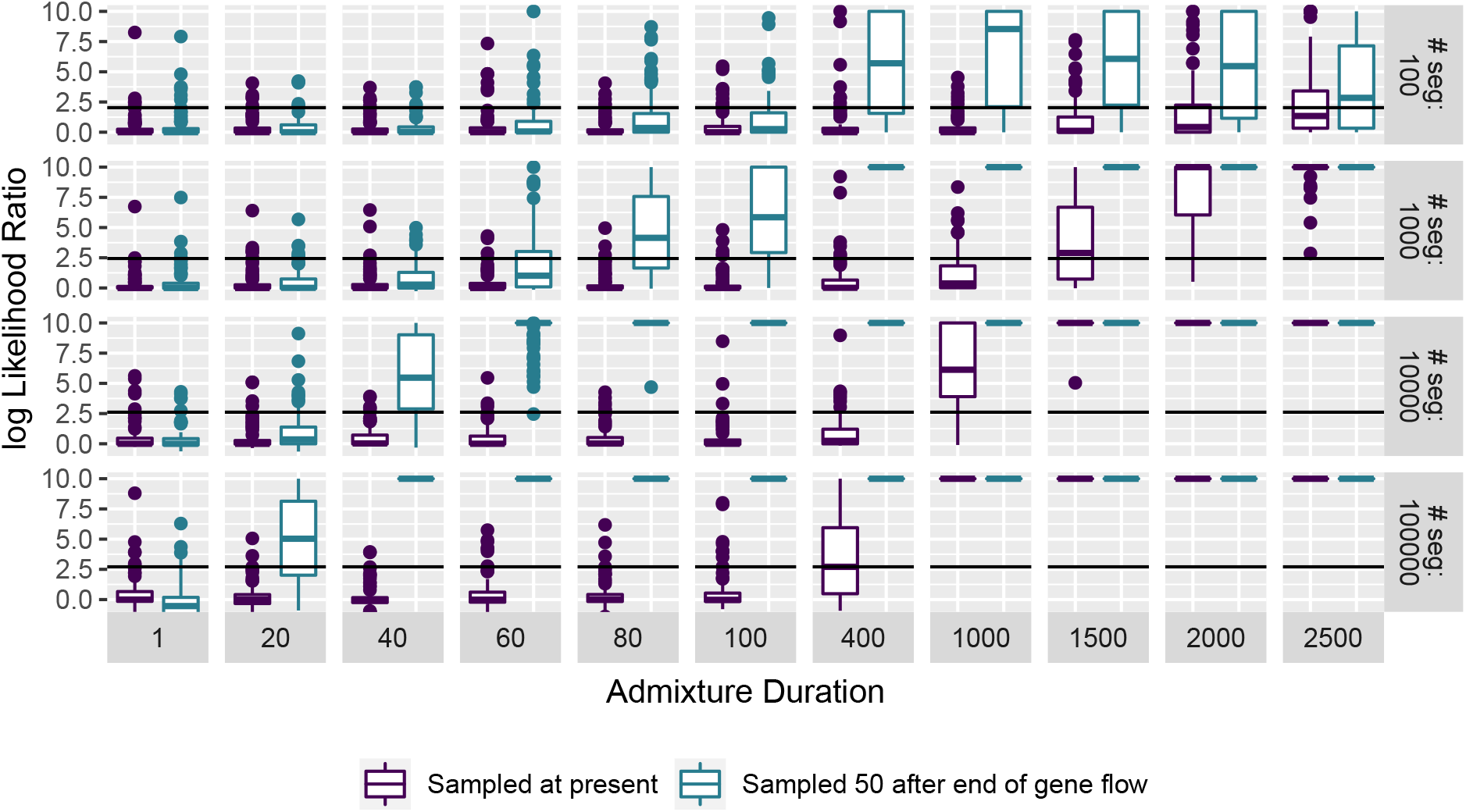
Model comparisons on perfect data. Segments are either sampled at the present (purple) or 50 generations after the end of gene flow (turquoise). Log-likelihood ratios bigger than ten are rounded to ten.

### Population Genetic Model Comparisons

In the previous section, we have shown that we can distinguish long pulses from instantenous gene flow under idealized conditions. As a more realistic scenario, we perform population genetic simulations using `msprime` (Kelleher et al., 2016). Throughout, we simulate 3% Neandertal admixture into non-Africans using a demographic model of archaic introgression (Supplementary Figure 1B) with a mean admixture time of 1,500 generations ago and varying durations. We simulate 20 chromosomes of length 150MB, using either a constant recombination map, or using the HapMap recombination map (HapMap Consortium, 2007). This results in ~ 10,000 introgressed segments. We then perform inference either using the simulated segments, segments inferred from the data (Skov et al., 2018), or ALD calculated using ALDER (Loh et al., 2013). We further vary recombination rate settings as i) inference and simulation under constant recombination rate (Constant/Constant), ii) simulation using the HapMap genetic map (HapMap Consortium, 2007), and inference using no correction (HapMap/Constant), iii) simulation using HapMap, correction using a different map (HapMap/AAMap) (Hinch et al., 2011), iv) and inference using the same map used for the simulations (HapMap/HapMap).

Using these simulations, we perform model comparisons (Figure 3A). For segments, we again use the likelihood-ratio and find that the results for the simulated segments closely match the simulations under the model (Figure 2), showing that our model is a good approximation in the parameter range of interest. In contrast, we find that for inferred segments, results greatly depend on the recombination rate used: For a constant recombination rate, results are similar, but for the HapMap-recombination map, we do not have any power to distinguish these scenarios. As we fit ALD using non-linear least squares, no formal model-comparison framework exists. Qualitatively, we plot the normalized residual sum-of squares (RSS) and find that they increase with *t_d_* for both recombination scenarios, suggesting that the difference between the two models increase.

**Figure 3:**
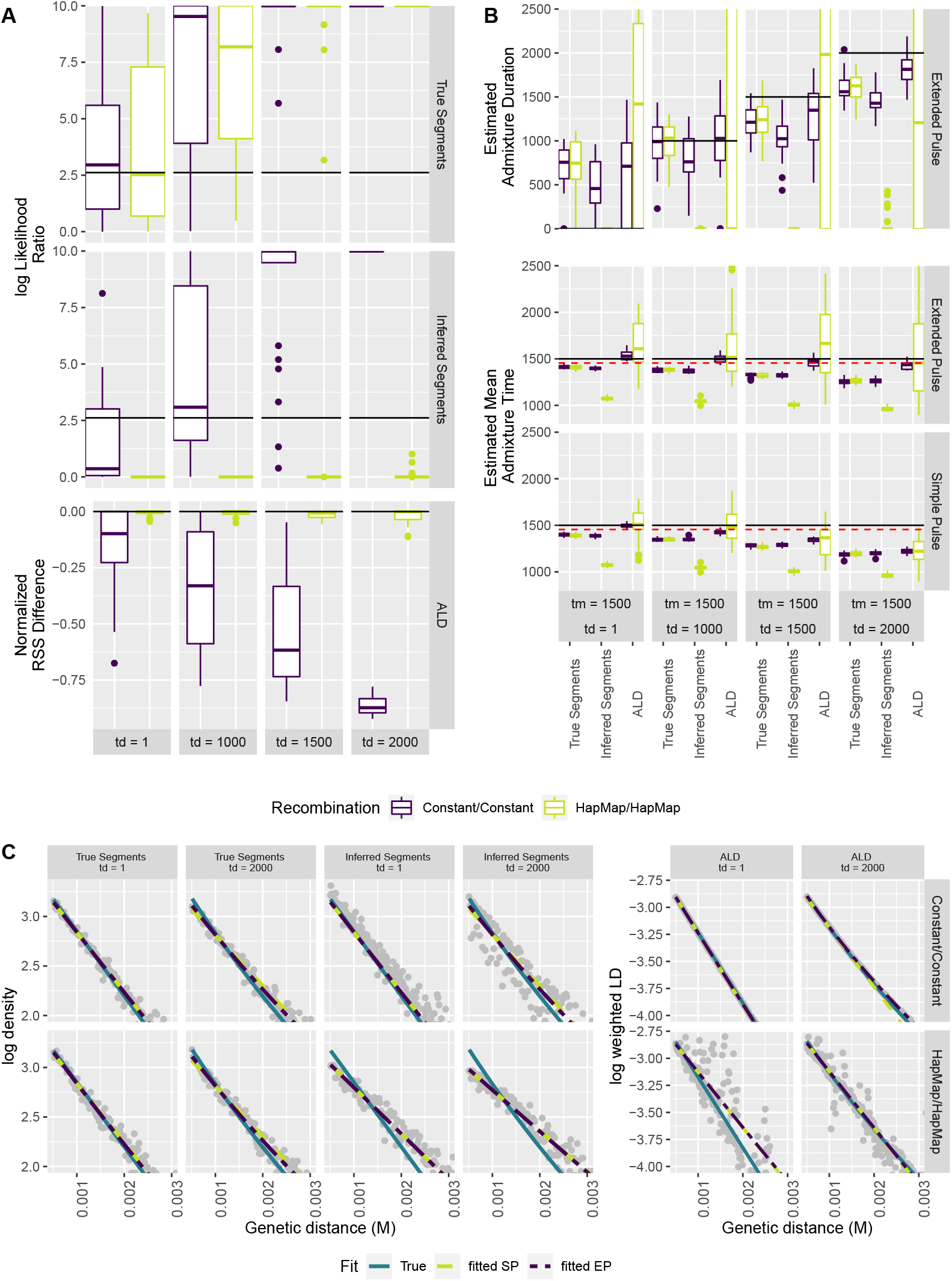
Model Choice, Model Fit and Parameter Estimates. A) Log-likelihood ratios and RSS difference between the simple and extended pulse models for segment data and ALD, respectively. Simulation and inference were done using constant(purple) and an empirical (teal) recombination map. B) Estimates of *t_m_* and *t_d_*. Solid black line indicate simulation values, the red dotted line adds a migration corrected (*t_m_*(1 − *α*)). C) Model fit for a single simulation in each scenario. Estimated segment density or weighted LD (grey) is compared with the expected (turquoise) and fitted single pulse (yellow) and extended pulse (purple).

Next, we evaluate parameter inference. In Figure 3B, we present estimates of the mean admixture times, admixture duration and the fitted segment and ALD distributions, respectively. We find that the mean admixture times are reasonably accurately estimated in most scenarios, the exception being the inferred segments when using the variable (HapMap) recombination map. The admixture duration estimates are often less accurate, and in most cases has very large variation between simulations.

We detect a slight, but consistent underestimate of the mean admixture times, which increases with *t_d_*. For the segments, this underestimate is likely due to the slight downward bias caused by recombination and coalescence between admixed segments (Liang and Nielsen, 2014, see also Appendix Genetic Drift and Recombination). For ALD, this bias is much less severe, particularly for inference under the extended pulse model. For scenarios where the recombination map is misspecified, *t_m_* is estimated to be only around half of its true value (Supplementary Figure 2). However, we find that in some cases, the extended pulse model provides a better estimate of *t_m_* by estimating the pulses to be extremely long.

In Figure 3C we show examples of the estimated segment length and ALD distributions compared to the simulated data. For these log-plots, the slope of the curve corresponds to the estimate of *t_m_*, and the deviation from linearity reflects the duration of gene flow. In all cases, we find that the expected decay is very close to linear, matching our finding that power to differentiate these old events is limited. We find that particularly when using a constant recombination map, all three summaries give a very close fit, and the segment length and ALD-decay distribution closely follow their expectations, which is consistent with the generally good parameter estimates under these conditions. In the case of a variable recombination map, we find that particularly inferred fragments perform poorly, which is reflected by a substantial downward bias of *t_m_* and *t_d_*.

### Comparing Effect Sizes for Technical Covariates

As we find that ALD performs as good or better than inferred segments (Figure 3), we focus on ALD for the remainder of this paper. Our next goal is to more carefully evaluate the relative importance of common assumptions made in the inference of admixture times, under both the simple and extended pulse model in the ALD framework on the bias and accuracy of estimates of *t_m_* under either model.

In particular, we use a Bayesian Generalized Linear Model (GLM) framework to contrast the effect of extended gene flow on admixture time inference with i) the effects of a simple/complex demographic history (Supplementary Figure 1), ii) recombination map variation, iii) the ALD ascertainment scheme, iv), *d_0_*, the minimum genetic distance between variants and v) the number of makers used to estimate the ALD curve (see Methods for details). For each modelling parameter and gene flow model, we use a simple model as the base case, and we study the impact of a more “realistic” alternative model.

In Figure 4, we present the estimated effect sizes for these six variables and four key interaction terms. To model bias, we fit a model to the standardized difference between the true and estimated mean admixture time, and to model accuracy (Methods, Supplementary Table 1, Supplementary Figure 3, Supplementary Table 2). These effect sizes are estimated using simulations under all possible parameter combinations on a scenario with admixture happening 1,500 generations ago. (Supplementary Figures 4 and 5).

**Figure 4:**
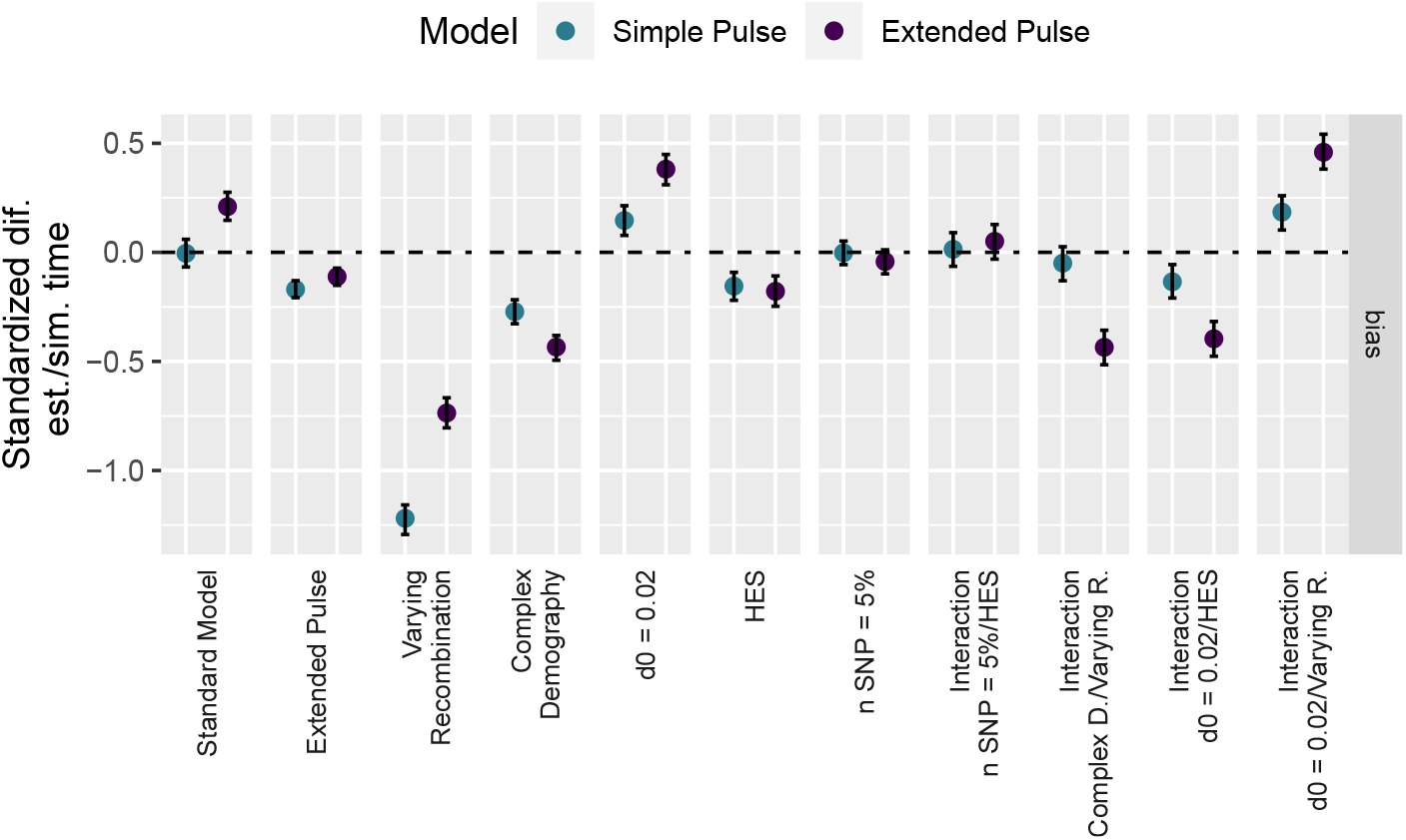
GLM effect sizes for the bias between simulated and estimated mean admixture time and 95% C.I. for the parameters between the simple and extended pulse model: gene flow (simple/extended), recombination rate (constant/varying), demography (simple/complex), minimal genetic distance (0.02/0.05 cM), SNPs used for ALD calculation (100 % / 5 %) and ascertainment scheme (LES/HES). Estimates are calculated across all possible combinations of parameters. Dotted horizontal line indicates unbiased admixture estimates.

As a baseline, for comparison, we define a standard model using a simple demography (Supplementary Figure 1A) and a constant recombination rate. This baseline model results in unbiased estimates of *t_m_* under the single pulse model with low deviation of 0.08 (0.02 – 0.14 *C.I*._95%_), and a slight upward bias 0.21 (0.15 – 0.28) for the extended pulse model.

The effect of simulating an extended-pulse gene flow only results in a slight bias of −0.17; −0.21 – −0.13) for the simple pulse and no bias for inference under the extended pulse model (−0.11; −0.15 – 0.07). In contrast, uncertainty in the genetic map causes by far the largest downward bias (Simple pulse: −1.22, −1.29 – −1.16; Extended Pulse: −0.74, −0.80 – −0.67) with high deviation in the estimates (Supplementary Figure 3). The more complex demography results in an underestimate of *t_m_*, presumably because of increased genetic drift, for both the simple pulse (−0.27, −0.33 – 0.22 and extended pulse models (−0.43, −0.49 – −0.38). The remaining parameters largely only have very minor effects, the biggest of which is changing the minimum cutoff from 0.05cM to to 0.02 cM.

### Application to Neandertal Data

Our next aim is to apply our model on the case of Neandertal gene flow into Eurasians. We estimate the Neandertal admixture pulse from the 1000 genomes data (The 1000 Genomes Project Consortium, 2015) and three high-coverage Neandertal genomes (Prüfer et al., 2013, 2017; Mafessoni et al., 2020) by fiting pulses with durations ranging from one generation up to 2,500 generations to the ALD decay curve (Figure 5, Supplementary Table 3). Plotting these best-fit ALD curves shows the extremely slight difference predicted under these drastically different gene flow scenarios (Figure 5A). The difference between scenarios becomes more apparent if we log-transform the y-axis (Figure 5B), where we see that ongoing gene flow results in a heavier tail in the ALD distributions. However, these LD values are very close to zero, and are thus only very noisily estimated.

**Figure 5:**
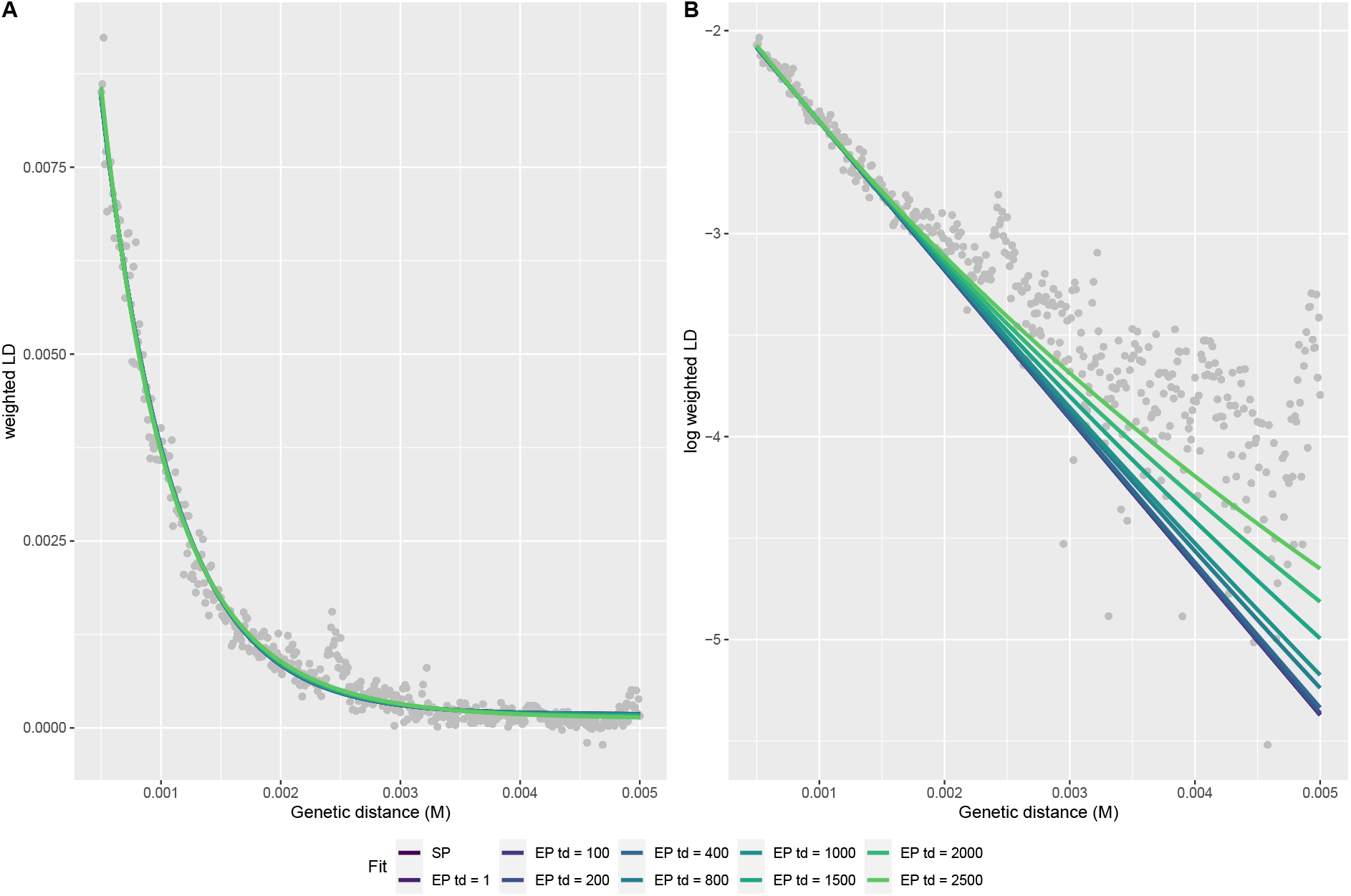
Neandertal Gene Flow in Modern Humans. We fit models with fixed *t_d_* from one to 2,500 generations of gene flow to the ALD-curve calculated from CEU individuals on A) natural and B) log-scale.

For short gene flows (less than 1,000 generations), our estimates for *t_m_* are very similar and identical to the simple pulse, at around 1,682 (1,526 – 1,839 *C.I*._95%_) generations. Extremely high values of *t_d_* result in slightly higher values of *t_m_* with overlapping compatibility intervals; but all predict that Neandertals would have survived until 30kya, for which the archaeological evidence is extremely sparse (Hublin, 2017). From the residual sum-of-squares, the models perform equally well, with longer extended pulses of gene flow achieving marginally better fits (Supplementary Table 4). Therefore, we find that all scenarios are compatible with the observed data, and that there is little power to differentiate these cases from genetics alone.

### Sampling Closer to the Admixture Event

Since Neandertal gene flow happened long in the past, much of the signal has been lost, and we have shown that in this scenario, power to distinguish different scenarios is low.

However, we have also shown in Figure 2 that inference is easier for more recent gene flow, a case that is relevant for many study systems. We investigate this in a series of simulations where the time between sampling and gene flow is smaller (Figure 6). We use the simple demographic scenario with a constant-sized populations (Supplementary Figure 1), and use ALD for inference using the optimised settings for the Neandertal case (ascertainment scheme = LES and d0 = 0.05 cM).

**Figure 6:**
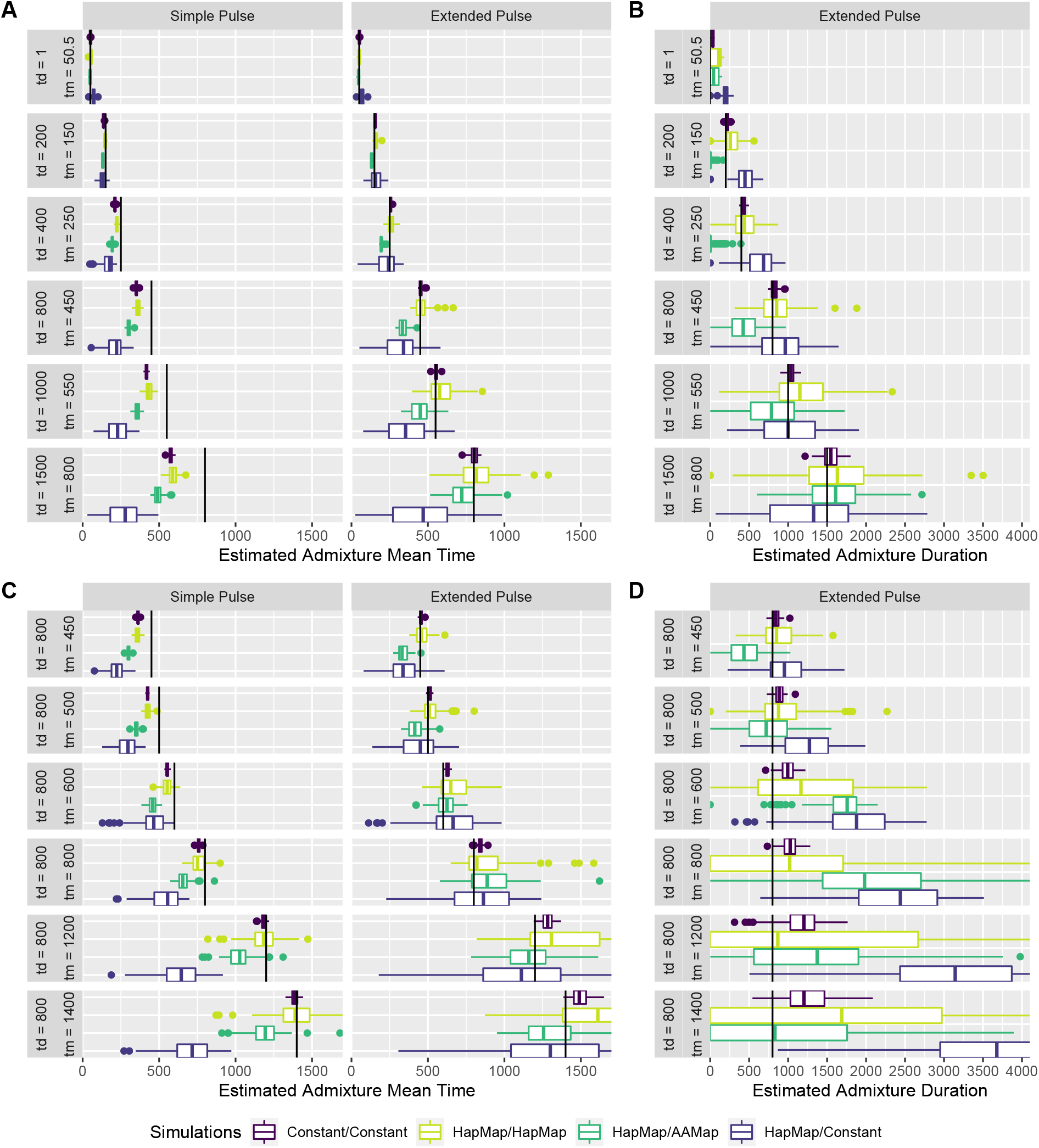
Parameter estimation using ALD from recent admixture pulses. We show estimates of *t_m_* (panels A, C) and *t_d_* for a series of simulations with increasing *t_d_* and *t_m_* such that we sample 50 generations after gene flow (top) and a series of simulations with a increasingly older extended pulse (bottom). All times are given in generations.

In Figure 6A and B, we show the accuracy of estimating *t_d_* and *t_m_* for increasingly longer pulses, sampled 50 generations after gene flow ended. The corresponding comparison of model fit is depicted in Supplementary Figure 6. For these cases, we find that inference of *t_m_* under the simple pulse model works well for the shortest pulses, but becomes increasingly downward biased as *t_d_* increases. Estimates of *t_m_* are less biased for inference under the extended pulse model, where we get accurate estimates particularly if recombination is constant. In the scenarios with a variable recombination rate, we find that for short, recent pulses, all corrections give good results, but for longer pulses, particularly assuming a constant recombination rate leads to a stronger bias. We also find that we are able to accurately infer the admixture duration, particularly if the recombination rate is constant.

In 6 C and D, we keep the pulse duration constant at *t_d_* = 800, but move it successively further into the past. For the first two cases of *t_m_* = 450 and *t_m_* = 500 where the pulse is recent, we again obtain good parameter estimates, but performance deteriorates for *t_m_* ≥ 600, which also results in low power to distinguish the simple and extended pulse model (Supplementary Figure 6, lower panel).

## Discussion

In this paper, we introduce a new population genetic model for dating extended pulses of gene flow. Our model has just two parameters, that can be interpreted as the mean time and duration of gene flow; and has simple closed form solutions for the segment length and ALD distributions. We show that both the instantaneous pulse and constant migration models are special cases of our model, where the duration is extremely short or long, respectively. We also demonstrate that the segment length distribution and ALD-decay can be directly transformed into each other; in particular the segment-length distribution is proportional to the second derivative of the ALD-decay curve. This makes our theory and models generally applicable beyond gene flow between Neandertals and humans. In fact, we find that we have little resolution for the parameter settings relevant for archaic gene flow, as the data resulting from simple and extended pulses long in the past is extremely similar. In contrast, we have much more power to estimate the duration of gene flow from events in the recent past, a scenario relevant for many hybridizing species. One limitation of our approach is that we assume that the overall amount of introduced material is low, and that we ignore the effects of genetic drift and selection.

Previous approaches to date Neandertal-human gene flow have focused almost entirely on the mean time of gene flow using a simple pulse model, for which reasonably tight credible intervals can be estimated (Sankararaman et al., 2012; Moorjani et al., 2016). Under this model, the credible intervals of this time are bounds of when gene flow between Neandertals and early modern humans could have happened.

Our estimate of the *t_m_* for Neandertal gene flow of 1,682 generations corresponds to a mean time estimate of 49ky (assuming a generation time of 29 years Moorjani et al. (2016)), with bounds of 44-54ky. This is in almost perfect agreement with the previous result of (Moorjani et al., 2016), (41-54ky), which is based on largely the same method. However, here we show that models of extended gene flow with *t_d_* up to a thousand generations provide very similar fits to the data; and that marginally better fits are achieved with very long gene flows. However, these models all would have Neandertals survive until around 30kya, whereas archaeological evidence for Neandertals surviving beyond 40ky is increasingly sparse (Hublin, 2017), so that these models of extremely long gene flow might be rejected on these grounds.

Our finding that the observed data is compatible with models involving hundreds of generations of gene flow means that while likely substantial amounts of gene flow happened around these mean times, gene flow might have also happened tens of thousands of years before or after. This is of great practical importance, as it makes linking genetic admixture date estimates with biogeographical events much more difficult (Sankararaman et al., 2012; Lazaridis et al., 2016; Douka et al., 2019; Jacobs et al., 2019; Vyas and Mulligan, 2019).

The discovery of early modern human genomes dated to 40,000 − 45,000 ya with very recent Neandertal ancestors less than ten generations ago (Fu et al., 2014; Hajdinjak et al., 2021) illustrates that gene flow likely happened over at least several thousand years. In general, inference based on ancient genomes (Fu et al., 2014, 2015; Moorjani et al., 2016; Hajdinjak et al., 2021) promises to resolve some of these dating issues, as inference is substantially easier when admixture is more recent, as the time difference between gene flow and sampling time is much lower (Figures 2, 6). However, using these genomes for dating leads to further hurdles, particularly pertaining to the spatial distribution of admixture events; whereas we can assume that the spatial structure present in initial upper paleolithic modern humans is largely homogenized in present-day people, the introgression signals observed in Bacho-Kiro and Oase could be partially private to these populations, and thus these populations may have a different admixture time distribution than present-day people.

The uncertainty over the duration of Neandertal gene flow also has some implications for selection on introgressed Neandertal haplotypes. Neandertal alleles have been suggested to be deleterious in modern human populations due to an increased mutation load (Harris and Nielsen, 2016; Juric et al., 2016). Some details of these models may be affected if migration occurred over a longer time. For example, Harris and Nielsen (2016) suggested that an initial pulse of gene flow of up to 10% Neandertal ancestry might be necessary to explain current amounts of Neandertal ancestry, with very high variance in the first few generations after gene flow. More gradual gene flow could mean that such high admixture proportions were never reached, but rather a continuous migration-selection balance process persisted for the contact period, where deleterious Neandertal alleles continually entered the modern human populations, but were selected against immediately. However, in terms of the overall frequencies, there is likely little difference. For example, Juric et al. (2016) showed using a two-locus model that the frequencies of Neandertal haplotypes alone cannot be used to distinguish different admixture histories.

In addition, we find that modelling and method assumptions have an impact on admixture time estimates that are of a similar or larger magnitude than the effect of assuming a one-generation pulse. In particular, recombination rate variation poses a practical limitation to the accuracy of admixture date estimates for old gene flow, and has to be very carefully considered when making inferences about admixture times. A possible reason is that both an extended pulse, as well as a non-homogeneous recombination map, lead to an admixture segment distribution that deviates from the expected exponential distribution. Throughout, we measure segment lengths and LD-decay distance in recombination units. Misspecification of the recombination rate will increase the variance in ALD or segment lengths, which might be confounded with a longer admixture pulse (Sankararaman et al., 2012). Therefore, population-specific fine-scale recombination maps are needed for accurate admixture time estimates, at least for admixture that happened more than a thousand generations ago. Estimates of more recent admixture appear to be more robust, perhaps because coarser-scale recombination maps are better estimated, differ less between populations (Hinch et al., 2011) and the error relative to fragment length is substantially lower.

To further refine admixture time estimates, time series data from more admixed early modern human and Neandertal genomes are needed. In particular, measures based on population differentiation (e.g Wall et al., 2013; Browning et al., 2018; Villanea and Schraiber, 2019) hold much promise to understand the different events that contributed to archaic ancestry in modern humans. While Neandertal ancestry in present-day people has been largely homogenized due to the substantial gene flow between populations, samples from both the Neandertal and early modern human populations immediately involved with the gene flow could refine when and where this gene flow happened.

## Material & Methods

### Power Analysis under the Model

To test the power to distinguish the simple from an extended pulse we simulated 100, 1000, 10,000 and 100,000 unique times *T_i_* form a Gamma distribution, with shape parameter *k* + 1 and scale *k/t_m_*, setting *t_m_* to 1,500 generations. Segment lengths *L_i_* are obtained by sampling for each *T_i_* from an exponential distribution with rate parameter *T_i_* for present day samples and 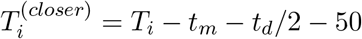 for sampling 50 generations after the end of gene flow. We obtain maximum-likelihood estimates for the simple (Eq. 13a) and extended pulse (Eq. 17) using the optim function implemented in R (R Core Team, 2019).

### Coalescent Simulations

We further test our approach on coalescence simulations using `msprime` (Kelleher et al., 2016). We focus on scenarios mimicking Neandertal admixture, and choose sample sizes to reflect those available from the 1000 Genomes data (The 1000 Genomes Project Consortium, 2015). For ALD simulations, we simulate 176 diploid African individuals and 170 diploid non-Africans, corresponding to the number of Yoruba (YRI) and Central Europeans from Utah (CEU). For inference based on segments we simulated 50 diploid non-Africans. Since three high coverage Neandertal genomes are available (Prüfer et al., 2013, 2017; Mafessoni et al., 2020) we simulate three diploid Neandertal genomes.

The demographic parameters are based on previous studies dating Neandertal admixture (Sankararaman et al., 2012; Fu et al., 2014; Moorjani et al., 2016; Skov et al., 2018). In the “simple” demographic model (Supplementary Figure 1 A), the effective population size is assumed constant at *N_e_* = 10, 000 for all populations, the split time between modern humans and Neandertals is 10,000 generations and the split between Africans and non-Africans is 2,550 generations. The migration rate from Neandertals into non-Africans was set to zero before the split from Africans, to ensure that there is no Neandertal ancestry in Africans. For a more complex scenario of human population history, we followed the demographic model simulation setup of Skov et al. (2018), but only simulated the Europeans. The split time of non-Africans are kept the same as in the ALD simulations (2550 generations ago) (Supplementary Figure 1B).

For each individual we simulate 20 chromosomes with a length of 150 Mb each. The mutation rates are set to 2 * 10^−8^ and 1.2 * 10^−8^ per base per generation for the “simple” and “complex” models, respectively. The recombination rates are set to 1 * 10^−8^ per base pair per generation for the simple demography and 1.2 * 10^−8^ per base pair per generation for the complex demographic model, unless specified otherwise.

Since inferring archaic segments is slow, we use 25 replicates for scenarios where we compare segment-based and ALD-based inference, and use 100 replicates when we only perform ALD-based inference.

#### Simulating Admixture

We specify simulations under the extended pulse model using the mean admixture time *t_m_* and the duration *t_d_*. We recover the simple pulse model by setting *t_d_* = 1, up to errors due to discrete generations. To obtain the migration rates in each generation, we use a discretized version of the migration density (Eq. 15b), which we then scale to the approximate amount of Neandertal ancestry in non-Africans (*α* = 0.03).

#### Recombination Maps

Uncertainties in the recombination map were previously shown to influence admixture time estimates (Sankararaman et al., 2012; Fu et al., 2014; Sankararaman et al., 2016). To investigate the effect of more realistic recombination rate variation, we perform simulations using empirical recombination maps. For the GLM, we use the African-American map (Hinch et al., 2011) for simulations and for the remaining simulations we use the HapMap phase 3 map (HapMap Consortium, 2007). For simplicity, we use the same recombination map (150 Mb of chromosome 1, excluding the first and last 10 Mb) for all simulated chromosomes. When simulating under an empirical map, with the analysis assuming a constant rate (*i.e*. no correction), we use the mean recombination rate from the respective map to calculate the genetic distance from the physical distance for each SNP. The mean recombination rate is calculated from the 150 Mb map (1.017 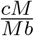 AAMap, 0.992 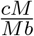 HapMap). For inference, each segment is either assigned a lengths based on its physical length (“constant”), the African-American map or HapMap recombination map, depending on the inference scenario.

### Estimating Admixture Time from Simulated Segment Data

For estimating admixture time and duration from introgressed segments, we either used the simulated segment lengths directly, or alternatively added an inference step using the HMM from Skov et al. (2018). We only considered inferred segments with an average posterior probability of 0.9 or higher. Furthermore, we use an upper and lower cutoff for inferred segment length of 0.05 cM and 1.2 cM. We fit the simple (Eq. 13a) and extended pulse (Eq. 17) using the `optim` function implemented using `R 4.0.3` (method=“L-BFGS-B”) with lower and upper constrains being 1, 5000 for *t_m_* and 2,1*e*8 for *k*, respectively.

### Estimating Admixture Time from ALD Data

#### Ascertainment Scheme

Since ALD for ancient admixture events can be quite similar to the genomic background, SNPs need to be ascertained to enrich for Neandertal informative sites in the test population. This removes noise and amplifies the ALD signal (Sankararaman et al., 2012). We evaluate the impact of the ascertainment scheme by contrasting two distinct schemes (Sankararaman et al., 2012; Fu et al., 2014). The lower-enrichment ascertainment scheme (LES) only considers sites that are fixed for the ancestral state in Africans and polymorphic or fixed derived in Neandertals. The higher-enrichment ascertainment scheme (HES) is more restrictive in that it further excludes all sites that are not polymorphic in non-Africans.

#### ALD Calculation

The pairwise weighted LD between the ascertained SNPs a certain genetic distance *d* apart is calculated using ALDER (Loh et al., 2013). A minimal genetic distance *d*_0_ between SNPs is set either to 0.02 cM and 0.05 cM. This minimal distance cutoff removes extremely short-range LD, which might also be due to inheritance of segments from the ancestral population (incomplete lineage sorting ILS) and not gene flow.

#### Parameter Estimates

We estimate parameters by fitting the ALD-curve to equations 22 and 23 using a non-linear least square approach implemented in the `nls` function in `R 4.0.3` (algorithm=“port”) with lower and upper constrains being 1 and 5000 for *t_m_* and 1/1*e*10 and 1/2 for 1/*k*, respectively. To achieve better conversion we pre-fit the functions using the estimates of the `DEoptim` optimization (Ardia et al., 2016) as starting parameters for the `nls` function. To improve estimates for *t_d_*, we run the fitting using 10 iterations to avoid local optima. We select the model with the lowest residual-sum-of-squares.

### Modeling Parameter Effect Sizes

To estimate the effect size of the different parameters (Eq. 24) we use a Bayesian Generalized Linear Model, where *E* is the response, and *A, M, D, R, S* and *G* are binary predictors.

The model can be written as

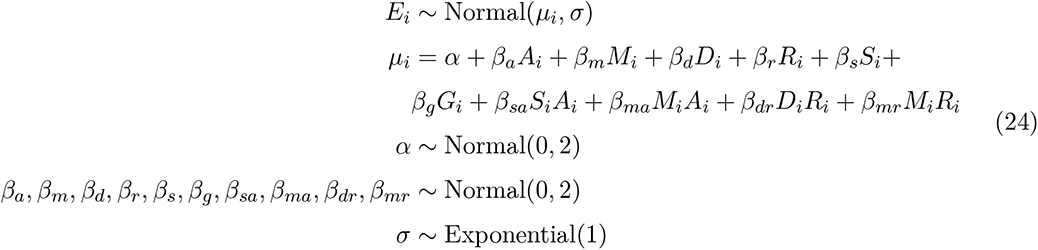

where the variables define

- ascertainment scheme: *A_i_* = LES/HES
- minimal genetic distance: *M_i_* = 0.02*cM*/0.05*cM*
- demography: *D_i_* = simple/complex
- recombination rate: *R_i_* = constant/variable
- n SNPs used: *S_i_* = 100 %/5 %
- gene flow model: *G_i_* = simple pulse (SP)/extended pulse (EP)

We fit two models, a primary model aimed at investigating the bias of our estimates, and a second model aimed at investigating the deviation. In the first case, the response variable *E_i_* is

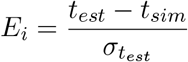
and in the second case we use the absolute error

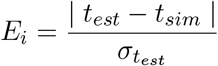
where *σ* is the standard deviation of *t_est_*. We also modeled the interaction between number of used SNPs and the ascertainment scheme (*β_sa_*), minimal distance and ascertainment (*β_ma_*), demography and recombination (*β_dr_*) and minimal distance and recombination(*β_mr_*).

We perform simulations using all possible parameter combinations. For the effect of the amount of SNPs i.e. accuracy of the ALD estimates, we downsampled the data by randomly choosing 5 % of the overall SNPs for ALD calculation. We define a standard model having a constant recombination rate, simple demography and gene flow, LES ascertainment and *d*_0_ = 0.05. The genetic distance is assigned from the physical position using the average recombination rate of the African-American genetic map (*i.e*., assuming the recombination rate is constant over the simulated chromosome given by this value) for simulations under a variable recombination rate. For each of the possible sets of parameters, we simulate 100 replicates each and fit ALD decay curves. We excluded a small number of simulations for which either the simple pulse or extended pulse curve could not be estimated (87 out of 6,400).

We assume a Normal likelihood because it is the maximum entropy distribution in our case. We obtained the posterior probability using a Hamiltonian Monte Carlo MCMC algorithm, as implemented in STAN (Carpenter et al., 2017) using an R interface (Stan Development Team, 2018; McElreath, 2020). The Markov chains converged to the target distribution (Rhat = 1) and efficiently sampled from the posterior (Supplementary Table 1 and 2).

### Estimating Neandertal Admixture Time

We estimate the Neandertal admixture time distribution using ALD from the 1000 Genomes data (The 1000 Genomes Project Consortium, 2015), together with the Altai, Vindija and Chagyrskaya high coverage Neandertals. We include the 107 unrelated individuals from the YRI as representatives of unadmixed Africans and all CEU as admixed Europeans. We only consider biallelic sites, and determine the ancestral allele using the Chimpanzee reference genome (panTro4). We used the CEU specific fine-scale recombination map (Spence and Song, 2019) to convert the physical distance between sites into genetic distance.

## Acknowledgments

We thank Svante Pääbo, Janet Kelso, Fabrizio Mafessoni, Stéphane Peyrégne, Laurits Skov, Divyaratan Popli, Alba Bossom Mesa, Arev Sümer and Shai Carmi for helpful comments and discussions. This work was supported by the Max Planck Society and the European Research Council (grant number: 694707) to Svante Pääbo.

## Author Contributions

- Conceptualization (Design of study) – BMP
- Software – LNMI
- Methodology – lead: BMP – support: LNMI, HR
- Formal Analysis – LNMI
- Visualization – LNMI
- Data Curation – LNMI
- Writing (original draft preparation) – lead: LNMI – support: BMP
- Writing (review and editing) – input from all authors
- Supervision – BMP

## Data Availability

The simulation script, the analysis pipeline and the *Extended Admixture Pulse* model is available on GitHub: https://github.com/LeonardoIasi/Extended_Admixture_Pulse. No new data were generated for this study.

## Appendix: Derivation and Approximation Details

### Formal Motivation for ALD

In the main text, we motivate the ALD decay using the intuitive argument of “adding up” ALD introduced at different generations in the past. Here we present a more formal derivation: In one generation, LD changes due to LD-decay through recombination, and through the introduction of new ALD through migration.

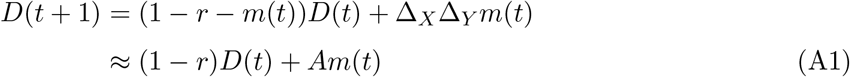

where *A* is a constant and *r* the map distance between the pair of markers. The approximation in the second line is valid if we ignore the effect of new introgressed material replacing older introgressed material.

This leads to the following linear differential equation:

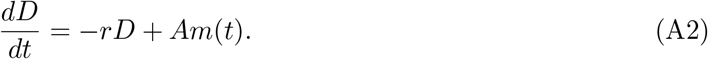

It is straightforward to verify that equation 6 is a solution to this equation:

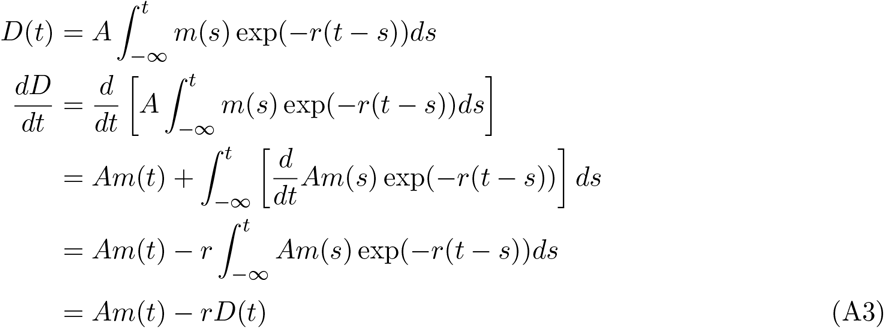

where the third line follows from Leibniz’ integral rule.

#### Replacement During Pulse

The “old”-LD also changes due to addition of new introgressed material, which is accomodated using

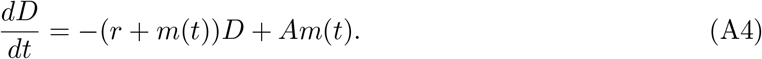

This equation has solution

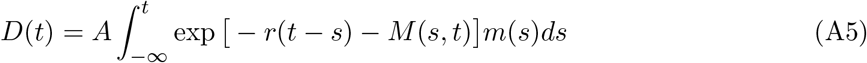

where

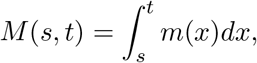

which can be interpreted as contributing how much LD has decayed from introgression time *s* to the observation time *t* due to replacement from new introgressed material.

This follows from

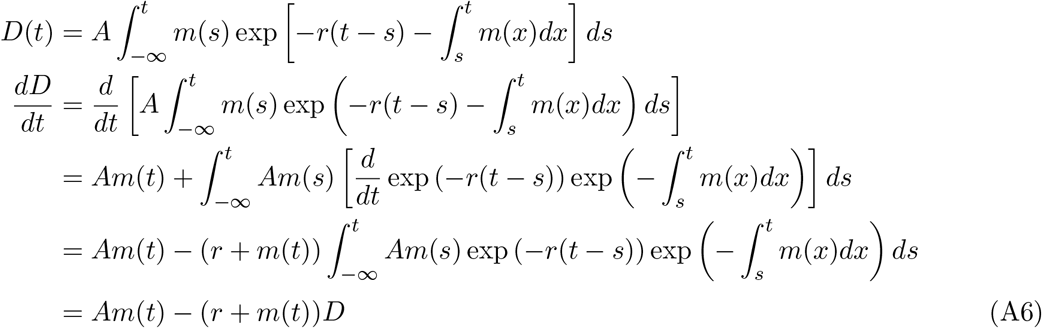

where the derivative can be evaluated using the product rule:

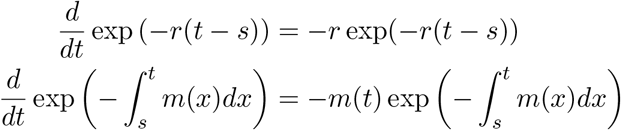

Changing the flow of time and setting *t* = 0 as in the main text, times, this results in

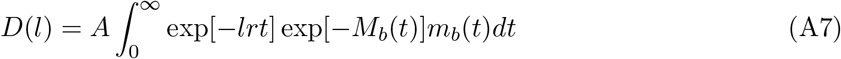

where again *m_b_*(*t*) = *m*(−*t*) and

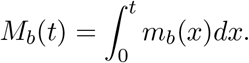

This motivates the “effective” migration rate

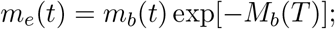

for the case of constant migration, *M_b_*(*t*) = *mt* and *m_b_*(*t*) = *me^−mt^*, which is an exponential density (Pool and Nielsen, 2009).

### Connection Between Admixture Segment Length Distribution and ALD Function

Here we describe how the admixture segment length distribution *P* (*l*) and the ALD function *D*(*l*) are interconnected (under the assumptions of this work) and in fact uniquely determine each other as claimed in the main text (Eq. 9 and Eq. 10).

Following the models outlined in the main text, throughout we assume that admixture is rare and that consequently admixture segments do not interact. Using these assumptions, we first describe how the admixture segment length distribution uniquely determines the ALD function, and second how the ALD function in reverse uniquely determines the admixture segment length distribution.

### From the Admixture Segment Length Distribution to the ALD Function

Without loss of generality, we assume that derived allele frequencies are 0 in the target and 1 in the admixture source population, otherwise the resulting ALD curve can be re-weighted with a constant factor *A* as in Eq. 4.

We start by noting that two-point ALD between two loci can be written as:

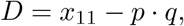

where *x*_11_ denotes the frequency of 11-haplotypes and *p* and *q* the allele frequencies at these two loci. Under the assumption that introgressed segments do not interact with each other, genome-wide excess 11 haplotypes beyond random association (*p* · *q* for each pair of loci) originate from pairs of markers on the same introgressed segments. To get the genome-wide average *D* we therefore have to sum over the contribution of 11 haplotypes from introgressed segments of all lengths.

For all pairs of markers a map distance *l* apart, only segments of length *x* > *l* contribute pairs of markers at distance *l*. Let us first describe a single introgressed segment of length *x*. In the limit of long chromosome (of length *G*) and of high marker density, a fraction (*x* − *l*)*/G* of all pairs of markers at distance *L* fall both onto this segment. Then, denoting the expected number of segments of length *x* as *E*(*x*), we sum over all segment lengths *x* > *l* to get:

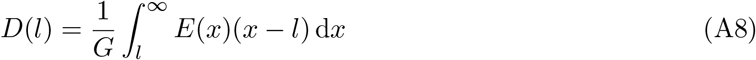

The expected number of segments of a given length *E*(*x*) can be directly derived from the segment length distribution via 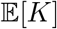, the total number of all introgressed segments:

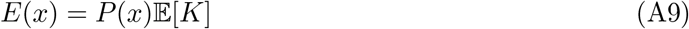

Plugging A9 into Eq. A8 yields:

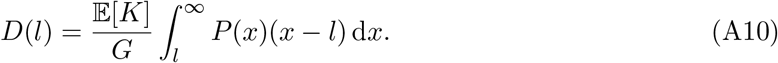

Eq. A10 now allows one to directly calculate *D*(*l*) from *P*(*l*), which shows that *P*(*l*) uniquely determines *D*(*l*).

### From the ALD Function to the Admixture Segment Length Distribution

To derive the inverse relationship, we start by differentiating *D*(*l*) twice with respect to 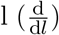. Using Eq. A10 and the Leibniz integral rule yields:

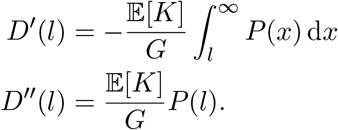

Simple rearrangement and plugging in the re-weighting of LD with *A* (that we omitted above) yields:

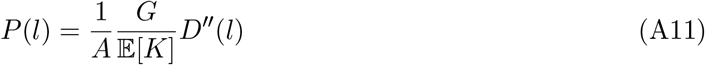

Thus, *D*(*l*) uniquely determines *P*(*l*).

### For the Continuous Admixture Model

For the concrete case of the continuous admixture model we derived explicit formulas for both the ALD function and the admixture segment length distribution (Eq. 3 and (Eq. 8). We can validate the above derived functional relationships (Eq. A10) directly and show that the more general derivation above holds for these specific formulas central for this work.

To check the relationships, we start with reiterating the explicit formulas: (Eq. 3) and (Eq. 8):

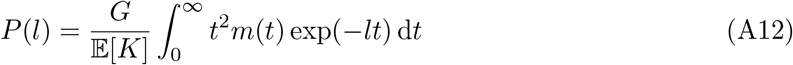

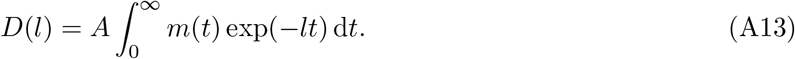

First, plugging Eq. A12 into the functional relationship Eq. A10 yields:

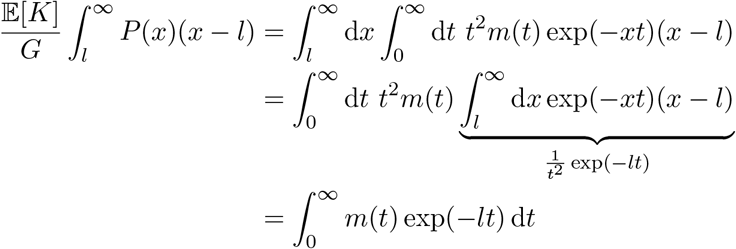

If we reweight the right-hand side with the allele-frequency differences A, it becomes D(l) from Eq. A13, which finishes the validation that the first functional relationship Eq. A10 holds.

To verify the second functional relationship (Eq. A11) we differentiate *D*(*l*) twice and multiply with 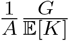:

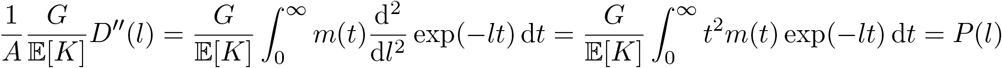

### Genetic Drift and Recombination

Our model assumes that Neandertal segments in the human population always recombine with non-Neandertal haplotypes, and that the effect of genetic drift can be neglected. In this appendix, we discuss some possible extensions of the model to incorporate aspects of genetic drift and recombination between admixture segments.

#### Single Pulse Theory of Recombination Between Fragments

Under our modelling assumptions, we show that 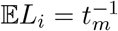, but this ignores recombination between introgressed material. Under a single pulse, Liang and Nielsen (2014) showed that under the SMC model (McVean and Cardin, 2005) the expectation is reduced as

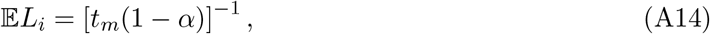

and under the SMC’ model (Marjoram and Wall, 2006) this is

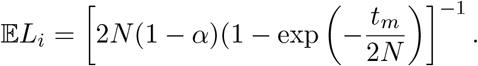

Using a Taylor expansion around *N* → ∞ and ignoring terms of the order 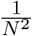, this can be approximated as

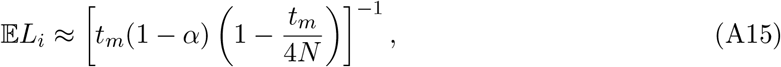

which makes the similarity to the SMC model more apparent.

This is compared to 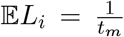 as we obtained from equation 14a. The (1 − *α*)-term models recombination between adjacent introgressed segments; and the 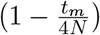 can be thought of reflecting genetic drift.

The justification for both of these formulas is that they are geometric mixtures of exponential distributions (Liang and Nielsen, 2014), which are themselves exponential. Under the extended pulse model, the segment length distribution is no longer exponential, so the segments may have a more complicated mixture distribution.

For the case of Neandertal admixture, assuming gene flow happened over a short duration, these equations can be used to estimate the error made from ignoring drift and recombination between Neandertal segments. As *α* ≈ 0.03, *t_m_* ≈ 1600, *N* ≈ 10, 000, and so the expected combined error of these two terms is on the order of 10%.

#### Effect of Reduced “Effective” Recombination and Coalescence

In the ALD framework, we can take further complications into account. For example, we can motivate

- an “effective” recombination rate *r*(*t*) = *r*(1 − *α*(*t*)) that takes into account as the admixture fraction increases, some recombination events will be between introgressed material, and we denote the total amount of introgressed material by time *t* as *α*(*t*).
- the allele frequencies in the admixing populations may change, so that we replace the constant *A* = Δ*_x_*Δ*_y_* by *A*(*t*) = Δ*x*(*t*)Δ*_y_*(*t*).
- genetic drift will fix some haplotypes, which then can no longer decay. This happens at rate 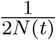

Taken together, the analogous equation is

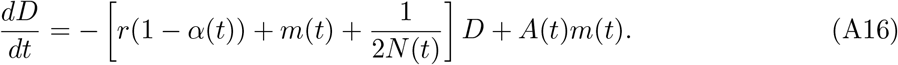

This equation is still a first-order non-homogeneous linear differential equation, so the solution will have the same form

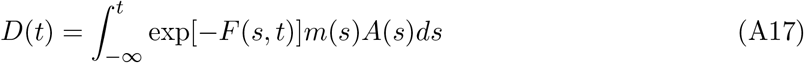

where

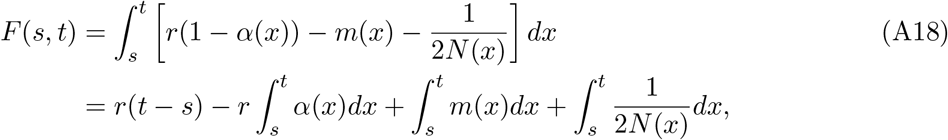

For example, if we assume *N*, *A*(*t*) and *α* are all constant, and migration is a simple pulse at *t_m_*

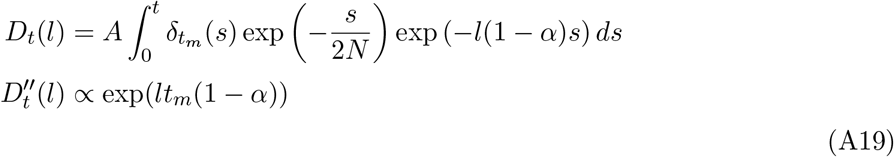

