## Supplement Material for "An extended admixture pulse model reveals the limitations to Human-Neandertal introgression dating"

### Supplement Figures

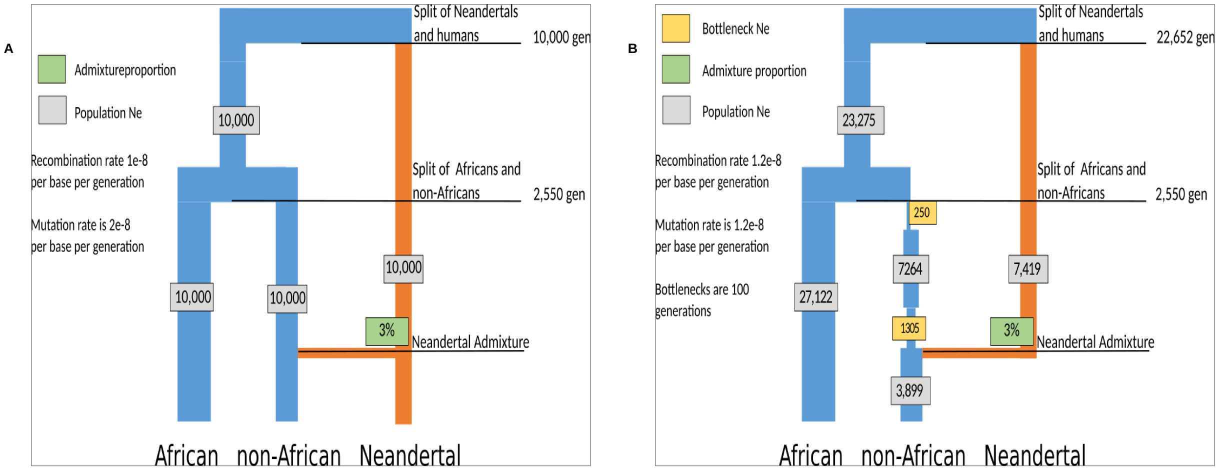


Figure 1: Demographic models of Neandertal introgression into non-Africans used for the simulations. A) Simple demographic model used for ALD simulations with constant population sizes. B) Complex demographic model with population size changes derived from Skov et al. 2018.

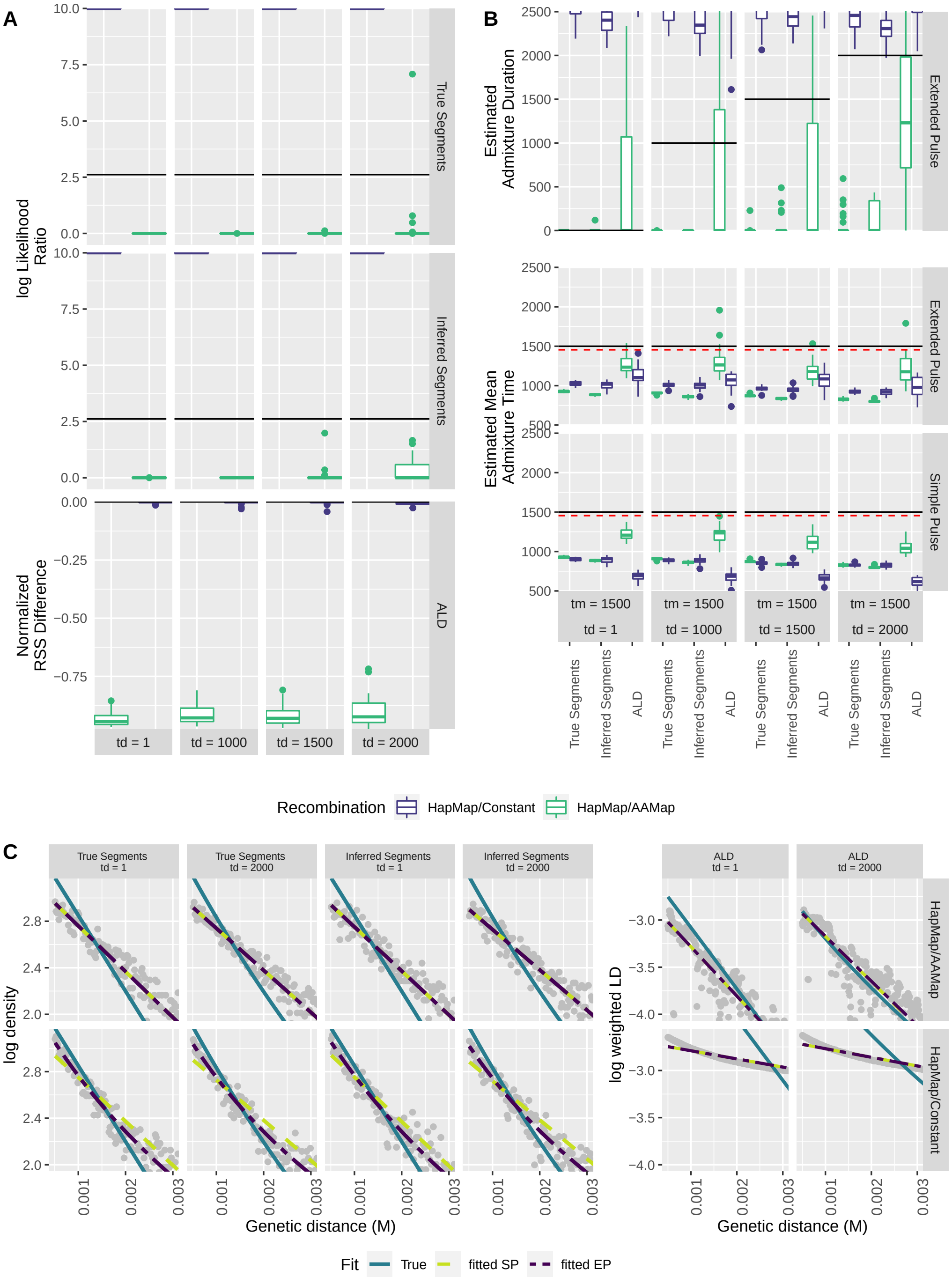


Figure 2: Comparison between the simple and extended pulse on true and inferred segment length and ALD decay estimates using an empirical recombination map (HapMap) for simulations with a fixed mean time ($t_m$) of 1,500 generations ago and varying durations ($t_d$). Genetic distances are assigned either using a constant rate (HapMap/Constant) or the AAMap (HapMap/AAMap). All times are given in generations. A) Log likelihood ratios between the two models for segment data and normalized difference between the residual sum-of-squares between the two models for ALD data. B) Mean time estimates of admixture and extended pulse estimate for admixture duration. Solid black line indicates true $t_m$ and $t_d$, red dotted line indicates migration corrected admixture time ($t_m$(1-m)). C) Comparison of the fit to data between the simple and extended pulse using true and estimated parameters.

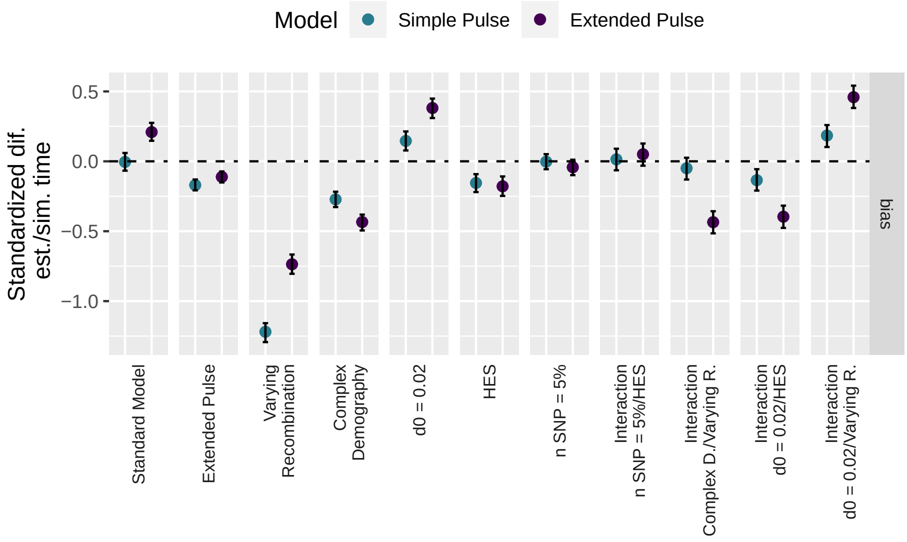


Figure 3: GLM effect sizes for the deviation between simulated and estimated mean admixture time and 95% C.I. for the parameters between the simple and extended pulse model: gene flow (simple/extended), recombination rate (constant/varying), demography (simple/complex), minimal genetic distance (0.02/0.05 cM), SNPs used for ALD calculation (100 % / 5 %) and ascertainment scheme (LES/HES). Estimates are calculated across all possible combinations of parameters. Dotted horizontal line indicates unbiased admixture estimates.

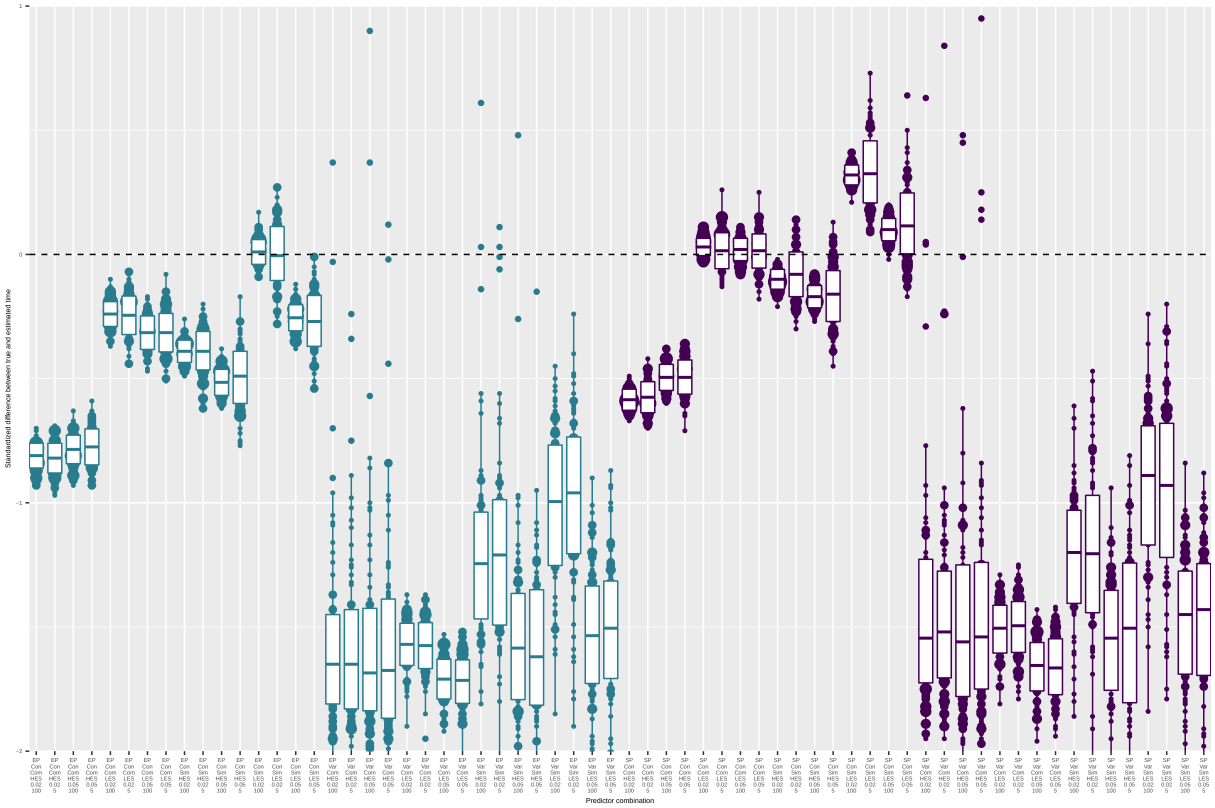


Figure 4: Comparison of the standardized difference between true and estimated mean admixture time under the simple pulse model for simulations of all combinations of parameters: ascertainment scheme = LES/HES, minimal genetic distance ($d_0$) = 0.02/0.05 cM, demography = simple/complex (sim/com), recombination = constant/variable (con/var), SNP used 100 % / 5 % and the gene flow model = simple pulse/extended pulse (SP/EP). Simulations with a simple pulse gene flow are indicated in purple, extended pulse in turquoise. Each simulation was repeated 100 times. Dotted horizontal line indicates no difference between true and estimated time.

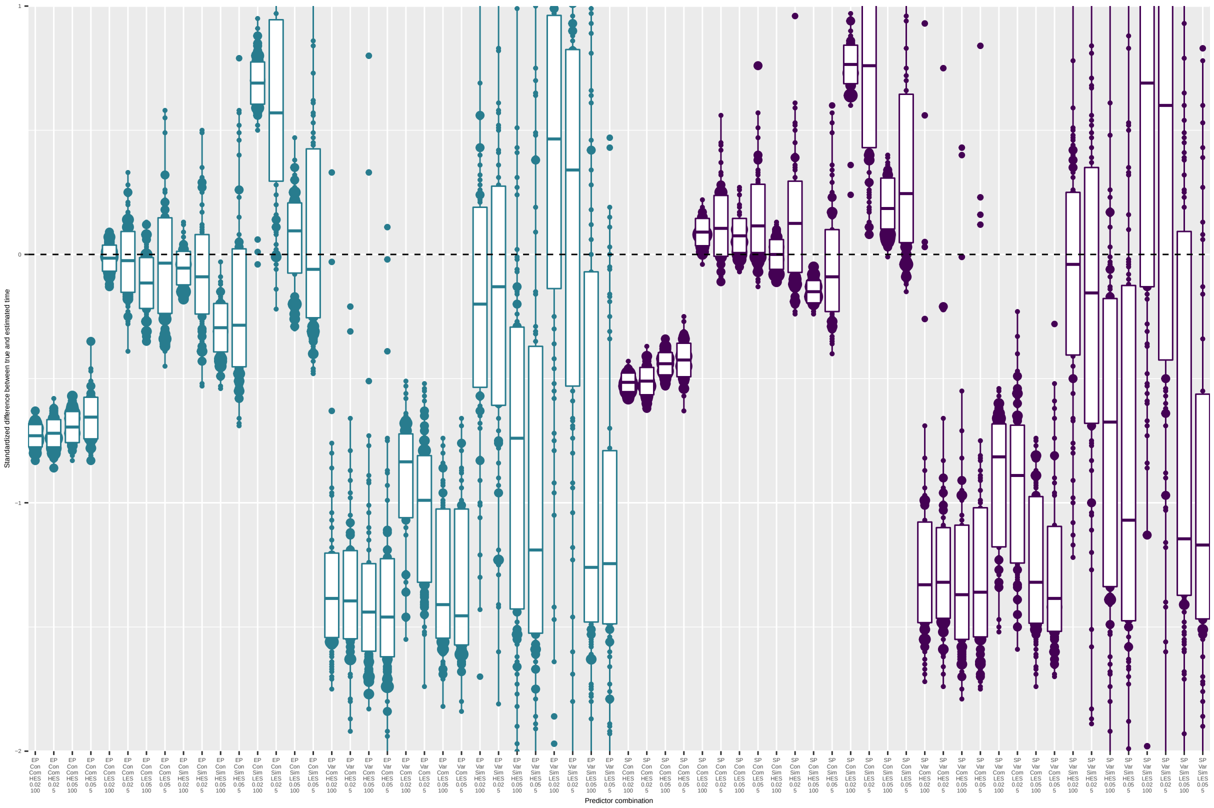


Figure 5: Comparison of the standardized difference between true and estimated mean admixture time under the extended pulse model for simulations of all combinations of parameters: ascertainment scheme = LES/HES, minimal genetic distance ($d_0$) = 0.02/0.05 cM, demography = simple/complex (sim/com), recombination = constant/variable (con/var), SNP used 100 % / 5 % and the gene flow = simple pulse/extended pulse (SP/EP). Simulations with a simple pulse gene flow are indicated in purple, extended pulse in turquoise. Each simulation was repeated 100 times. Dotted horizontal line indicates no difference between true and estimated time.

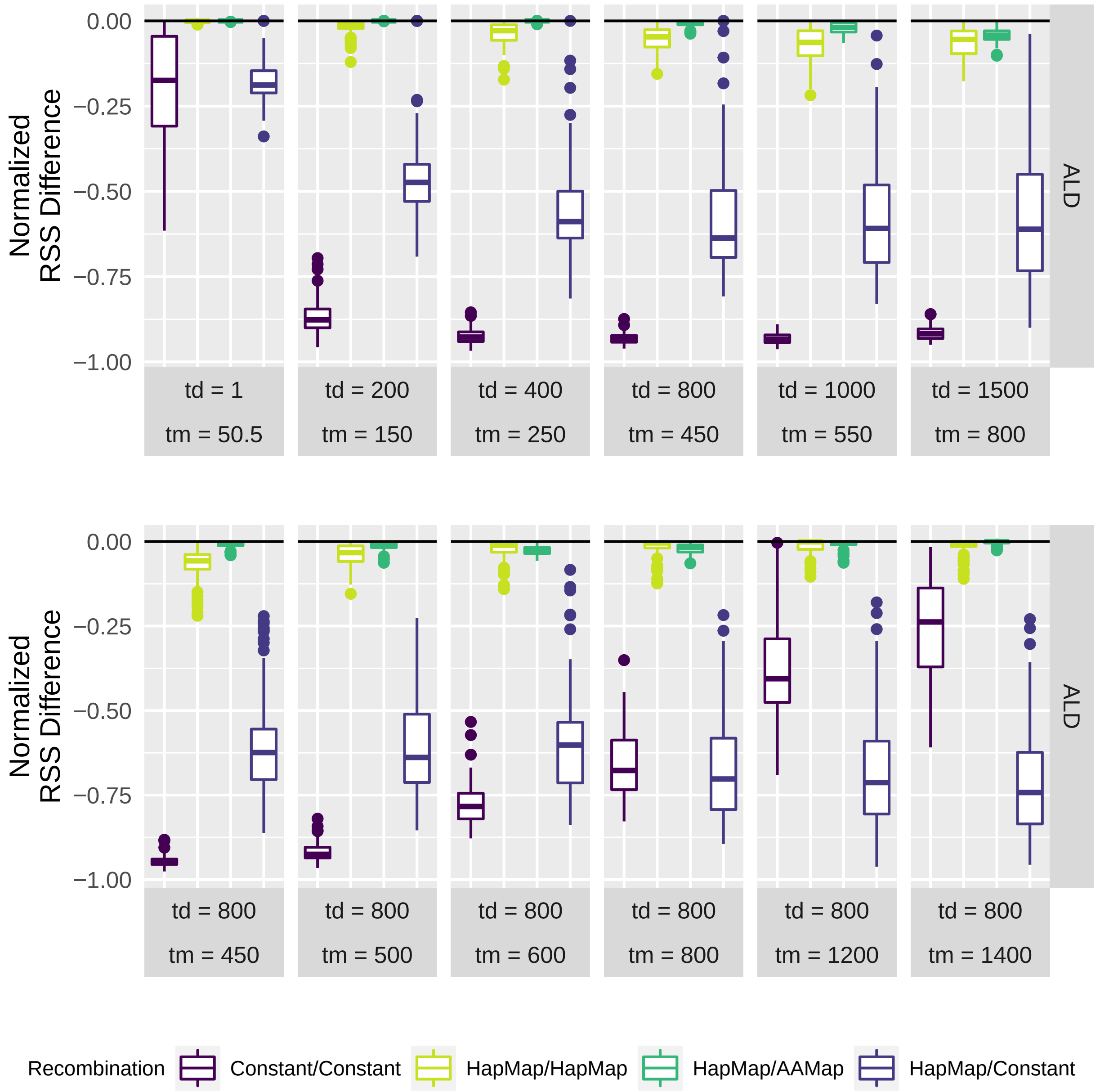


Figure 6: Sampling closer to the admixture event: Normalized difference in the residual-sum-of-squares between the simple pulse fit and extended pulse fit A) Comparison for scenarios sampled 50 generations after the gene flow ended, with different gene flow durations. B) Comparison for scenarios with different sampling times after 800 generations of gene flow.

#### Supplement Tables

Table 1: Comparing effect sizes for technical covariates: GLM parameter estimates on the bias between the true and estimated mean admixture time

| model | GLM | variable | mean | 5.5% | 94.5% | n eff | Rhat |
|---|---|---|---|---|---|---|---|
| Simple Pulse | bias | Standard Model | 0.00 | -0.07 | 0.06 | 1922 | 1 |
| | bias | Extended Pulse | -0.17 | -0.21 | -0.13 | 2439 | 1 |
| | bias | Varying Recombination | -1.22 | -1.29 | -1.16 | 2521 | 1 |
| | bias | Complex Demography | -0.27 | -0.33 | -0.22 | 2488 | 1 |
| | bias | d0 = 0.02 | 0.15 | 0.08 | 0.21 | 2282 | 1 |
| | bias | HES | -0.15 | -0.22 | -0.09 | 2507 | 1 |
| | bias | n SNP = 5% | 0.00 | -0.06 | 0.05 | 3171 | 1 |
| | bias | Interaction n SNP = 5%/HES | 0.01 | -0.06 | 0.09 | 2169 | 1 |
| | bias | Interaction Complex D./Varying R. | -0.05 | -0.13 | 0.03 | 2386 | 1 |
| | bias | Interaction d0 = 0.02/HES | -0.14 | -0.21 | -0.06 | 2032 | 1 |
| | bias | Interaction d0 = 0.02/Varying R. | 0.18 | 0.10 | 0.26 | 2086 | 1 |
| | bias | Sigma | 0.79 | 0.77 | 0.80 | 4267 | 1 |
| Extended Pulse tm | bias | Standard Model | 0.21 | 0.15 | 0.28 | 1876 | 1 |
| | bias | Extended Pulse | -0.11 | -0.15 | -0.07 | 2650 | 1 |
| | bias | Varying Recombination | -0.74 | -0.80 | -0.67 | 2284 | 1 |
| | bias | Complex Demography | -0.43 | -0.49 | -0.38 | 2678 | 1 |
| | bias | d0 = 0.02 | 0.38 | 0.31 | 0.45 | 2330 | 1 |
| | bias | HES | -0.18 | -0.25 | -0.11 | 2556 | 1 |
| | bias | n SNP = 5% | -0.04 | -0.10 | 0.01 | 3891 | 1 |
| | bias | Interaction n SNP = 5%/HES | 0.05 | -0.03 | 0.13 | 2196 | 1 |
| | bias | Interaction Complex D./Varying R. | -0.44 | -0.52 | -0.36 | 2375 | 1 |
| | bias | Interaction d0 = 0.02/HES | -0.40 | -0.48 | -0.32 | 2312 | 1 |
| | bias | Interaction d0 = 0.02/Varying R. | 0.46 | 0.38 | 0.54 | 2110 | 1 |
| | bias | Sigma | 0.81 | 0.79 | 0.82 | 4128 | 1 |

Table 2: Comparing effect sizes for technical covariates: GLM parameter estimates on absolute deviation between the true and estimated mean admixture time

| model | GLM | variable | mean | 5.5% | 94.5% | n eff | Rhat |
|---|---|---|---|---|---|---|---|
| Simple Pulse | abs. deviation | Standard Model | 0.08 | 0.02 | 0.14 | 1880 | 1 |
| | abs. deviation | Extended Pulse | 0.15 | 0.12 | 0.19 | 2456 | 1 |
| | abs. deviation | Varying Recombination | 1.26 | 1.19 | 1.32 | 3091 | 1 |
| | abs. deviation | Complex Demography | 0.17 | 0.12 | 0.22 | 2621 | 1 |
| | abs. deviation | d0 = 0.02 | -0.06 | -0.12 | 0.00 | 2293 | 1 |
| | abs. deviation | HES | 0.18 | 0.12 | 0.24 | 2347 | 1 |
| | abs. deviation | n SNP = 5% | 0.01 | -0.04 | 0.06 | 3529 | 1 |
| | abs. deviation | Interaction n SNP = 5%/HES | 0.00 | -0.07 | 0.07 | 2086 | 1 |
| | abs. deviation | Interaction Complex D./Varying R. | 0.03 | -0.03 | 0.11 | 2728 | 1 |
| | abs. deviation | Interaction d0 = 0.02/HES | 0.10 | 0.03 | 0.17 | 2336 | 1 |
| | abs. deviation | Interaction d0 = 0.02/Varying R. | -0.23 | -0.30 | -0.16 | 2206 | 1 |
| | abs. deviation | Sigma | 0.71 | 0.70 | 0.72 | 4341 | 1 |
| Extended Pulse tm | abs. deviation | Standard Model | 0.15 | 0.09 | 0.20 | 2138 | 1 |
| | abs. deviation | Extended Pulse | 0.09 | 0.05 | 0.12 | 3011 | 1 |
| | abs. deviation | Varying Recombination | 0.82 | 0.76 | 0.88 | 2697 | 1 |
| | abs. deviation | Complex Demography | 0.02 | -0.03 | 0.07 | 2665 | 1 |
| | abs. deviation | d0 = 0.02 | 0.11 | 0.05 | 0.17 | 2746 | 1 |
| | abs. deviation | HES | 0.10 | 0.04 | 0.16 | 2804 | 1 |
| | abs. deviation | n SNP = 5% | 0.07 | 0.02 | 0.12 | 4095 | 1 |
| | abs. deviation | Interaction n SNP = 5%/HES | 0.01 | -0.06 | 0.08 | 2698 | 1 |
| | abs. deviation | Interaction Complex D./Varying R. | 0.28 | 0.21 | 0.35 | 2705 | 1 |
| | abs. deviation | Interaction d0 = 0.02/HES | -0.03 | -0.10 | 0.04 | 2535 | 1 |
| | abs. deviation | Interaction d0 = 0.02/Varying R. | -0.41 | -0.48 | -0.34 | 2392 | 1 |
| | abs. deviation | Sigma | 0.71 | 0.70 | 0.72 | 4063 | 1 |

Table 3: Application to Neandertal data: Estimates

| | Model | tm | A | c |
|---|---|---|---|---|
| Simple Pulse | Simple Pulse | 1682 (1526 - 1839) | 0.019 (0.019 - 0.019) | 0.000182 (0.00016 - 0.000204) |
| Extended Pulse (td=1) | Extended Pulse (td=1) | 1682 (1526 - 1839) | 0.019 (0.019 - 0.019) | 0.000182 (0.00016 - 0.000204) |
| Extended Pulse (td=100) | Extended Pulse (td=100) | 1683 (1527 - 1839) | 0.019 (0.019 - 0.019) | 0.000182 (0.00016 - 0.000204) |
| Extended Pulse (td=200) | Extended Pulse (td=200) | 1685 (1528 - 1841) | 0.019 (0.019 - 0.019) | 0.000182 (0.00016 - 0.000204) |
| Extended Pulse (td=400) | Extended Pulse (td=400) | 1691 (1534 - 1848) | 0.019 (0.019 - 0.019) | 0.00018 (0.000158 - 0.000202) |
| Extended Pulse (td=800) | Extended Pulse (td=800) | 1717 (1558 - 1877) | 0.019 (0.019 - 0.019) | 0.000174 (0.000152 - 0.000196) |
| Extended Pulse (td=1000) | Extended Pulse (td=1000) | 1736 (1575 - 1898) | 0.02 (0.02 - 0.02) | 0.00017 (0.000148 - 0.000192) |
| Extended Pulse (td=1500) | Extended Pulse (td=1500) | 1800 (1633 - 1967) | 0.02 (0.02 - 0.02) | 0.000155 (0.000134 - 0.000177) |
| Extended Pulse (td=2000) | Extended Pulse (td=2000) | 1882 (1707 - 2057) | 0.021 (0.021 - 0.021) | 0.000138 (0.000116 - 0.000159) |
| Extended Pulse (td=2500) | Extended Pulse (td=2500) | 1978 (1794 - 2161) | 0.022 (0.022 - 0.022) | 0.000118 (9.6e-05 - 0.00014) |

Table 4: Application to Neandertal data: RSS

| Model | RSS |
|---|---|
| Simple Pulse | 2.53e-05 |
| Extended Pulse (td=1) | 2.53e-05 |
| Extended Pulse (td=100) | 2.53e-05 |
| Extended Pulse (td=200) | 2.53e-05 |
| Extended Pulse (td=400) | 2.53e-05 |
| Extended Pulse (td=800) | 2.51e-05 |
| Extended Pulse (td=1000) | 2.50e-05 |
| Extended Pulse (td=1500) | 2.47e-05 |
| Extended Pulse (td=2000) | 2.44e-05 |
| Extended Pulse (td=2500) | 2.42e-05 |
